# Optimizing seagrass planting arrangements for animal benefits in a multi-habitat restoration seascape

**DOI:** 10.1101/2025.09.16.676667

**Authors:** Michael Sievers, Christopher J. Brown, Jasmine A. Rasmussen, Benjamin Nielsen, Rune C. Steinfurth, Mogens R. Flindt, Timi L. Banke, Ben L. Gilby, Rod M. Connolly

## Abstract

Restoring lost and degraded ecosystems to enhance biodiversity and ecosystem services is a global priority, and animal responses to the restoration of habitats are a critical but undervalued component. Identifying the key drivers of animal colonization in restored habitats provides critical insights for restoration practitioners seeking to maximize ecological outcomes. When integrated into predictive frameworks and spatial decision- support tools, this knowledge becomes especially valuable for strategic planning, particularly in complex multi-habitat restoration projects where spatial configuration remains a crucial yet understudied dimension influencing ecosystem recovery trajectories. We collect and analyze animal data from one of the world’s largest multi- habitat coastal restoration systems in Denmark, comprising restored seagrass (*Zostera marina*), boulder reefs and mussel reefs. Using fine-scale spatial patterns in population abundances, we develop spatially explicit predictions across the seascape for various seagrass restoration scenarios and produce a series of optimizations, showing that it is practical to configure restoration to optimize biodiversity objectives, including those linked with fished species. Species-specific responses translated to variable outcomes across restoration scenarios and optimizations. While the optimal number and arrangement of restored patches varied depending on the target species or species group (e.g., fisheries species or seagrass specialists), one near-ubiquitous arrangement was patchy seagrass planting. This aligns with current practice, maximizes restoration efficiency, and highlights the importance of not homogenizing seascapes for biodiversity. Our approach provides a practical framework for incorporating animal monitoring data into restoration planning, helping practitioners design and optimize spatial planting configurations to achieve specific ecological objectives.

**Open Research Statement:** All data and code/scripts (R language), including a README file, are freely available at: https://github.com/msievers100/DenmarkSpatial

## Introduction

Ecosystem restoration is a global priority, driven by international initiatives such as the United Nations Decade on Ecosystem Restoration (2021–2030), the Kunming-Montreal Global Biodiversity Framework, and the EU Nature Restoration Law (Maron et al. 2018, Hering et al. 2023). These calls to action and legal frameworks highlight the urgent need to restore habitats to safeguard and enhance biodiversity and ecosystem services (Young and Schwartz 2019). Restoration not only aims to rebuild the structural component created by habitat-forming taxa (e.g., vegetation), it plays a crucial role in reestablishing animal populations that use and depend on these habitats (Gann et al. 2019, Sievers et al. 2024). These animals support many of the ecosystem services that restoration seeks to enhance and often perform ecological functions vital to ecosystem persistence and resilience, and thus restoration success (McAlpine et al. 2016, Reeves et al. 2020, Sievers et al. 2022). They can, however, also act as an impediment when their populations or behaviors disrupt desired ecological processes (Infantes et al. 2016, Sievers et al. 2022). Thus, understanding the factors that influence animal use of restored habitats can help guide restoration planning to enhance outcomes and meet objectives, particularly when this information is incorporated into predictive models and spatial optimizations (Gilby et al. 2018). Testing and optimizing the spatial arrangement of restored habitats is particularly important in a multi-habitat restoration context but remains a relatively unexplored facet in restoration science (Gilby et al. 2018, Vozzo et al. 2023).

Multi-habitat restoration – which involves the simultaneous or sequential restoration of more than one habitat type – offers a promising strategy for enhancing restoration outcomes, particularly for coastal marine systems (McAfee et al. 2022, Vozzo et al. 2023, Hackett et al. 2024). Since coastal seascapes are naturally heterogeneous, with documented daily, tidal, and ontogenetic movements of animals across different habitats, they are ideal systems for conducting and studying multi-habitat restoration approaches (Olds et al. 2016, Vozzo et al. 2023). Coastal habitats are also experiencing significant losses worldwide (Beck et al. 2009, Dunic et al. 2021) and are consequently the focus of accelerating restoration efforts (Duarte et al. 2020). Multi-habitat restoration benefits from facilitative interactions that occur among coastal ecosystem types and their inhabitants, and the premise that many animals use a mosaic of habitats during different life stages or to meet varying ecological needs (Pittman et al. 2021, Scherer-Lorenzen et al. 2022, Silliman et al. 2024, Vozzo et al. 2024). For instance, photosynthesizing aquatic plants can benefit from the restoration of filter-feeding animals such as mussels and oysters that increase water clarity (Wall et al. 2011), whilst many studies have demonstrated benefits for fishes and crustaceans of high connectivity among coastal ecosystems such as seagrass meadows, mangrove forests and coral reefs (Unsworth et al. 2008, Henderson et al. 2017, Olson et al. 2019). The strategic positioning of restored habitat patches relative to existing natural and other restored habitats can thus influence ecological outcomes. Yet, robust spatial models based on empirical data remain scarce for coastal restoration (Gilby et al. 2018).

Understanding the spatial factors that influence animal use of restored habitats is fundamental to developing predictive models and robust optimization models that can inform restoration (Boström et al. 2011, Gilby et al. 2018). The characteristics and arrangement of habitat patches – i.e., their size, shape, proximity to other habitat types, and positioning within the broader seascape – can profoundly affect colonization rates, species interactions, and ecosystem function recovery (Boström et al. 2006, Boström et al. 2011). Optimising site selection to meet successful restoration outcomes such as survivorship and proliferation of habitat forming species (Elliott et al. 2022) and socio- cultural objectives (Howie et al. 2024) have already been well documented in the literature. It is also likely important to utilize spatially explicit models when designing restoration projects to maximize outcomes for animals and supporting the facilitative interactions that can enhance both restoration success and economic outcomes. Predictive tools based on high-resolution spatial data and empirical monitoring can forecast how species respond to restoration interventions under different spatial configurations of restored habitats, allowing practitioners to compare multiple planting arrangements before implementation (Theuerkauf et al. 2019, Lester et al. 2020, Bertelli et al. 2022).

Such spatial optimization approaches are especially valuable for multi-habitat restoration, where the proximity and configuration of different habitat types influences species abundance, diversity, and ecosystem service provision (Vozzo et al. 2023, Vozzo et al. 2024). Although most studies on habitat proximity in coastal areas focus on scales >100s meters, facilitative interactions and effects of habitat configuration can also occur at finer scales (e.g., 1-10 meters; Vozzo et al. 2024), warranting data and models at finer resolution.

Here, we surveyed mobile fauna using a robust sampling arrangement within natural seagrass, on bare sand, and in restored seagrass at one of the world’s largest multi-habitat restoration seascapes. We calculate spatially explicit predictions of animal abundance – for both individual and multi-species objectives – for a series of hypothetical seagrass restoration scenarios with varying planting strategies and an optimized arrangement (Figure 2). Five scenarios focus first on *where* to restore, which offers potential practical and logistical benefits, while the optimization approach uses greedy heuristics to maximize outcomes irrespective of logistics. We thus showcase how animal monitoring data can be leveraged to develop spatially explicit predictions and optimize spatial arrangements for objective-dependent restoration planning and decision-making.

## Materials and Methods

### Overview

We evaluated how the spatial arrangement of natural and restored habitats in a Danish fjord influences the abundance of marine species. We then developed and compared five alternative restoration scenarios, along with an optimization algorithm, to predict the impact of different spatial configurations of seagrass restoration on both individual species and multi-species assemblages. This approach allowed us to identify and map optimal restoration arrangements that maximize ecological outcomes tailored to different objectives.

### Site description and fauna sampling

Surveys were carried out near Sellerup Strand in Vejle Fjord, Denmark (55.687, 9.681; Figure 1). Seagrass (*Zostera marina*) has been transplanted here annually since 2019 (∼3.7 ha at the time of surveys). In 2022 a restored boulder reef was created (∼2.2 ha), and in 2021 blue mussels (*Mytilus edulis*) were distributed either side of the boulder reef (8 ha, 320 tons; Figure 1). This restoration aimed to: (1) reestablish lost habitat types (as boulder-fishing has reduced the extent of hard substrate in the estuary, and seagrass extent has reduced significantly from historical levels; Timmermann et al. 2020), (2) reduce hydrodynamic wind-induced stress to support eelgrass restoration and establishment, and (3) increase biodiversity and provide refuge for fauna, especially fish (Flindt et al. 2024). For additional site information, see Banke et al. (2024). We surveyed mobile fauna in May 2023 at 132 sites that were within natural seagrass, on bare sand, and in restored seagrass that was transplanted 1, 2, 3, or 4 years ago (Figure 1). Vejle fjord represents a site of ongoing seagrass restoration; we thus focused on monitoring within natural and transplanted seagrass as well as bare areas within the seascape. Sites were between 1 and 2m in depth. Netting involved pushing a 60cm wide shrimp net (mesh size 2mm) along the substrate in a 3 m long transect twice (i.e., 3.6 m^2^ total area). All netting was conducted by a single person to avoid variability in technique. This provided a rapid assessment but likely missed some individuals of larger and more mobile taxa capable of detecting and avoiding the net. Between each transect, animals were transferred to a bucket with fresh seawater. Onshore, each individual animal was photographed, weighed, and identified to the lowest taxonomic level (i.e., species).

**Figure 1.**
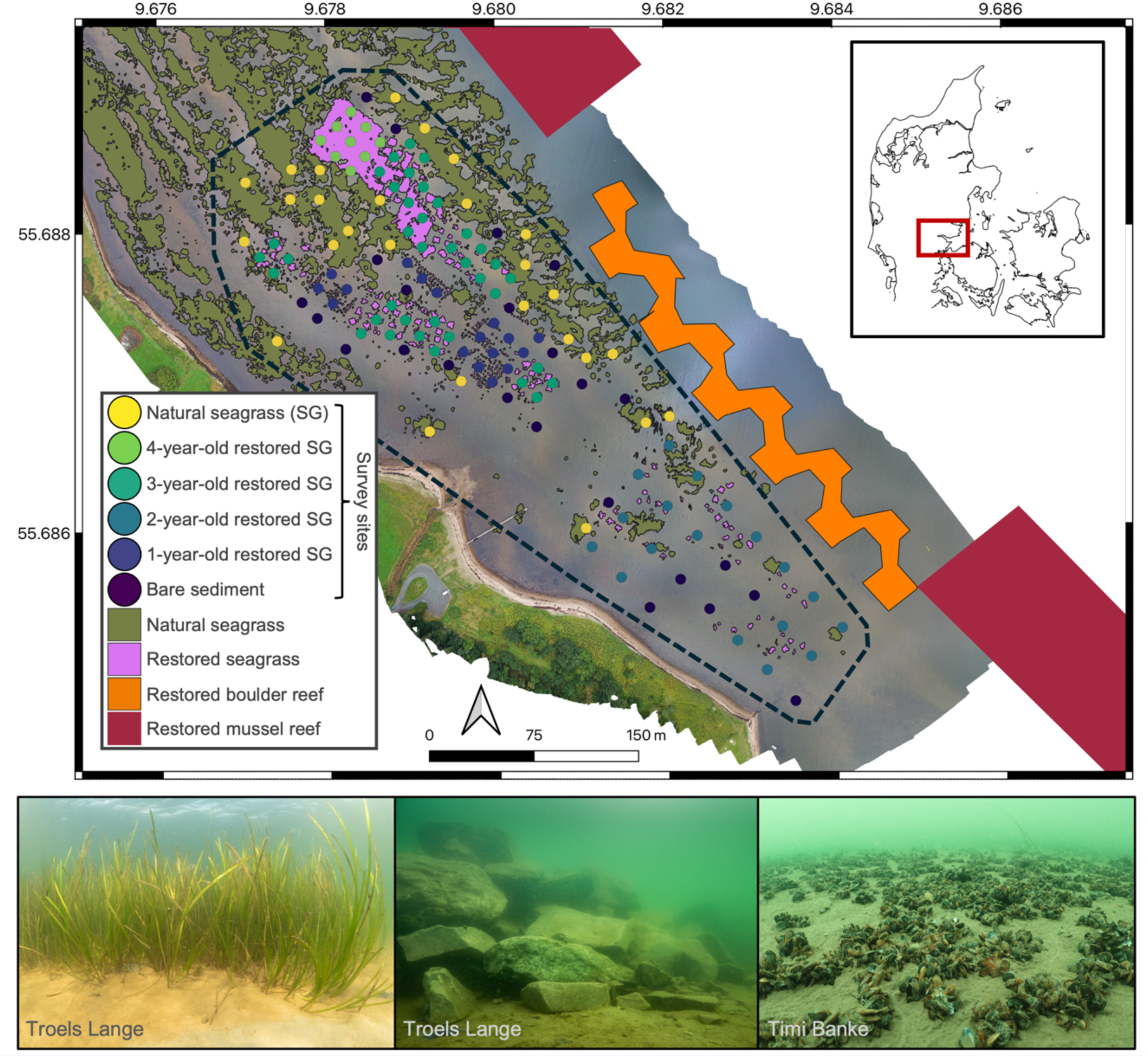
Visualization of the study site. Drone imagery of the study area in Vejle Fjord, Denmark, taken October 2023, showing the distribution of natural seagrass, restored seagrass, boulder reef and mussel reef, the location of the 132 survey sites, and the approximate region used for the seascape modelling (shown as a dashed-lined polygon). Below are images of the restored seagrass, boulder reef and mussel reef. We confirm permission for use in the manuscript of all photographs.

### Quantifying habitat and seascape characteristics

The area was mapped at 100m altitude using a drone (DJI Mavic 2 PRO) with a 20MP camera. Production of orthomosaics and Object-Based Image Analysis of seagrass extent was performed according to the procedure in Svane et al. 2021. An orthomosiac was produced with the software Agisoft Metashape (Ver. 1.7.3), while Object-Based Image Analysis was performed in Trimble eCognition Developer (Ver. 9.3) using a Support Vector Machine algorithm. Polygons for the boulder reef and mussel reef were created based on field-collected GPS waypoints. A series of seascape metrics were then calculated for each site using GIS software (QGIS Ver. 3.24), including: the area of seagrass (natural and restored) within a 10m radius, the perimeter to area ratio of seagrass (natural and restored) within a 10m radius, the Euclidian distance to the restored boulder reef, and Euclidian distance to the restored mussel reef. Given the spatial configuration of our sites, our focus on fine-scale interactions, and potential independence issues, we chose to focus on a radius of 10m to examine habitat-species relationships, and to develop and test our proof-of-concept approach.

### The influence of the arrangement of multiple restored and natural habitats on animals

We used Generalized Additive Mixed Models (GAMMs) in the *mgcv* package to evaluate species richness, Shannon diversity, and evenness with the fixed effects of: Treatment (categorical: bare, 1-, 2-, 3-, 4-year-old restored, and natural sites), the area of seagrass within a 10m radius (‘Area10m’), the ratio of perimeter to area of seagrass within a 10m radius (log-transformed; (‘RatioLog’), distance to the boulder reef (‘DistBouldEdge’), distance to the mussel reef (‘DistMussEdge’). Restored site maturity (i.e., time since restoration) can strongly influence species abundances (see Sievers et al. 2025); accounting for such factors allowed for the statistical parsing out of effects to evaluate the spatial effects more clearly. Continuous fixed effects had k set to 3. Site coordinates in UTM were included as a random effect (k = 4). All community-metric models used a Gaussian distribution. Species densities were modelled using the same model structure as community metrics, but with a negative binomial distribution. For all GAMMs, we used the *dredge* function in the *MuMIn* package to reduce full models (i.e., with all factors) to the best-fitting model (lowest AIC), and subsequently plot relationships between response and the remaining predictor variables within these reduced models. A distance matrix was constructed from UTM coordinates, and we examined both the spatial structure of residuals through variogram analysis and tested for spatial independence using Moran’s I statistic (Appendix S1: Table S1; all p-values >0.2). We checked for concurvity between model terms using the *concurvity* function in mgcv, which indicated no issues between model terms, except for between the spatial random effect and distance to boulder reef (Appendix S1: Table S2). Given that this reef is one continuous structure, this correlation is unsurprising. Finally, to investigate beta diversity, we examined the influence of model factors on species composition using distance-based redundancy analysis (dbRDA) in the *vegan* package. Species abundance data for the top 10 most abundant species were used to calculate a Bray-Curtis dissimilarity matrix. We then performed a permutation test with 999 iterations to assess the significance of the predictor variables in explaining variation in species composition.

We confirmed that our observational data had an experimental design that was valid for inferring the effects of multiple covariates and the restoration treatments. Density distribution plots revealed substantial overlap in distance-based covariates across all three habitat types, indicating that our sampling design achieved good spatial balance (Figure S1). This balance was further evidenced in the empirical cumulative distribution function (eCDF) plots, which show similar distributions between groups, particularly after matching (Figure S2). While Area10m showed expected differences among Treatment (with natural sites having higher seagrass coverage within 10m than restored sites, and bare sites having the least), the substantial overlap in spatial positioning relative to boulder and mussel reefs ensures that comparisons between habitat types are not confounded by systematic differences in proximity to these other restoration elements.

This covariate balance strengthens the inference from our statistical models by minimizing potential spatial biases that could arise in a purely opportunistic sampling design. Overall, our observation design was robust for statistically parsing out the spatial effects of habitat type and arrangement on animal communities.

### Predicting and mapping individual and multi-species abundance across restoration scenarios and optimizations

We developed nine objectives based on maximizing the individual abundance of lesser pipefish, common periwinkle, flatfish, whelks, Baltic prawns, and brown shrimp, and three multi-species groups: (1) all six species, (2) seagrass specialists (lesser pipefish, common periwinkle, Baltic prawns), and (3) fisheries species (flatfish, Baltic prawns). We scaled species abundances by dividing by the survey data standard deviations to ensure comparable contributions across species. For multi-species groups, the objective function was defined as maximizing the mean of scaled abundances across target species. To quantify the individual and multi-species abundances across the seascape, we used a negative binomial distribution with a pared down version of the GAMs, where the random spatial effect and the area to perimeter ratio were removed. The sampling design meant that the seascape predictors used in the GAMs were largely independent of each other, so we could partition different components of seascape effects on animal abundance.

We created a polygon around our sites (+10 m buffer), overlaid a grid to create a seascape of potential restoration sites that approximated the size of a typical transplantation patch within the fjord (N = 3973 ‘sites’ of 20.33m^2^). Potential restoration sites (N = 3300) included all bare areas and where current restored seagrass exists (i.e., restored seagrass was removed from the seascape), but excluded areas with natural seagrass, essentially providing a hypothetical seascape where no restoration had occurred.

We developed five exemplar restoration scenarios and an optimization algorithm to examine differences in objectives, related to where restoration took place (Figure 2). The scenarios were:

1. **Random site selection:** A null model that in reality is arguably a less feasible strategy due to the logistics of transplanting seagrass shoots, and a scenario that would not benefit from intra-specific facilitations benefitting transplant growth and survival (Valdez et al. 2020).
2. **Start-near-seagrass:** Start transplanting nearest to the bulk of the natural seagrass in the north-west of the seascape and radiating out, maximizing benefits from intra-specific facilitations and making animal movement from natural patches easier.
3. **Start-away-from-seagrass:** Start transplanting farthest away from the bulk of the natural seagrass in the south-east of the seascape and radiating out, maximizing the area of the seascape in which at least some seagrass occurs.
4. **Start-near-mussel-reef:** Start transplanting nearest to the mussel reef and radiating out, maximizing inter-habitat facilitative processes and animal movement.
5. **Start-near-boulder-reef:** Start transplanting nearest to the boulder reef and radiating out, maximizing inter-habitat facilitative processes and animal movement.

For these five scenarios, outcomes were calculated at each of 20 even ‘steps’, whereby bare sites converted to the equivalent of ‘4-year-old restored sites’ until the entire seascape transitioned into seagrass (containing both restored and natural). Since intraspecific facilitatory processes suggests transplanting close to other transplants can enhance seagrass success (Valdez et al. 2020), and the fact that restoring within smaller areas is less logistically challenging, we largely test scenarios that follow this logic (scenarios 2–5). At each step, models were updated iteratively to reflect changes in seagrass area in a 10m radius around sites (a dynamic covariate that changes as nearby sites transition from bare to seagrass). For the random site selection ‘null’ scenario, sensitivity was examined by calculating population abundance for 10 sets of random site selection.

**Figure 2.**
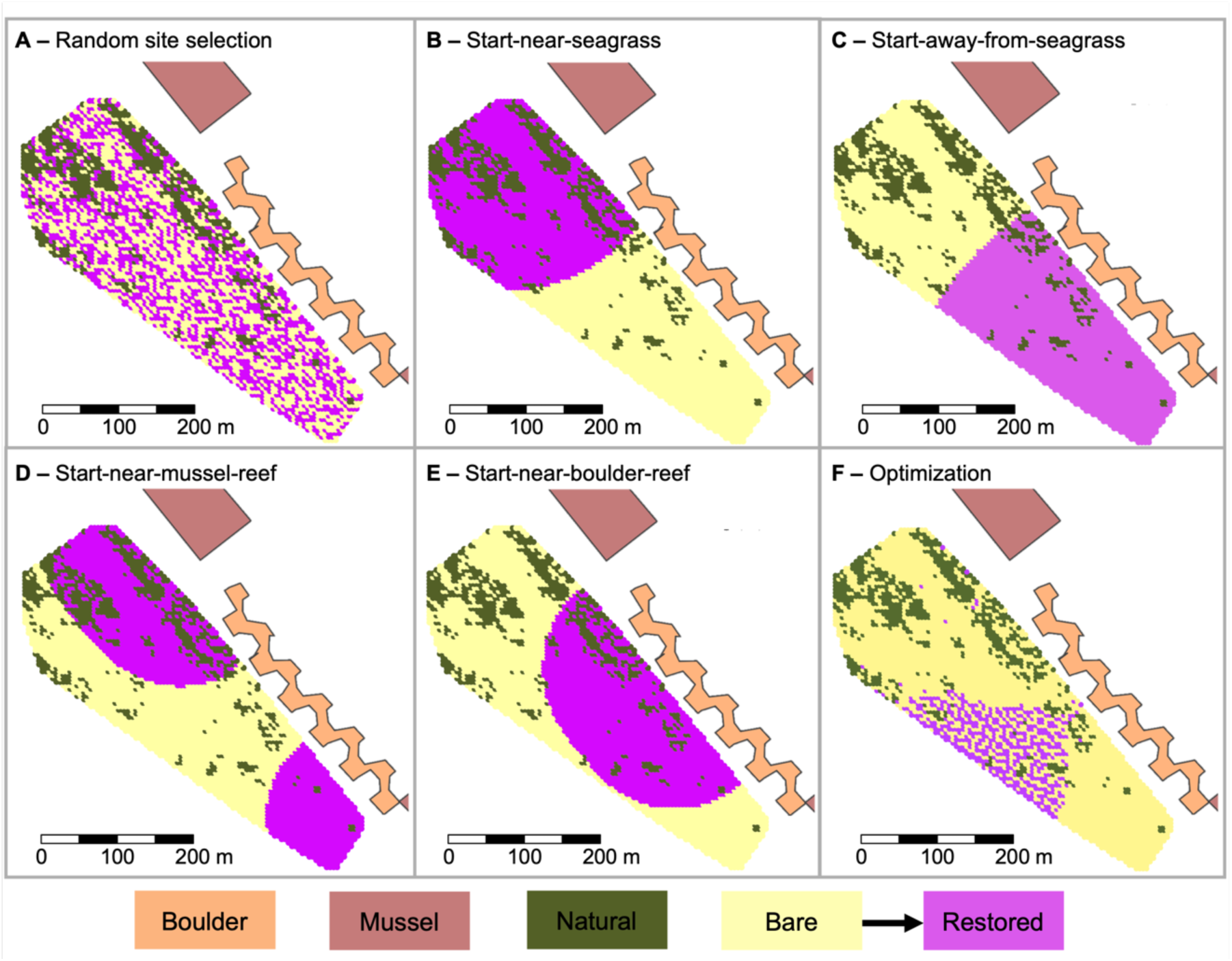
Visualization of the restoration scenarios and spatial optimization. The distribution of natural seagrass (olive) restored seagrass (pink), and bare (light yellow) sites at the halfway point (i.e., 50% of the bare seascape restored into seagrass) for (A-E) five scenarios, and a (F) theoretical optimal arrangement. The boulder reef (orange) and mussel reefs (salmon) are also shown.

For the optimal arrangement, we developed a spatially explicit greedy optimization algorithm to identify the configuration of restoration sites that would maximize individual and multi-species abundance (our objective function), based on:

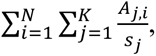

Subject to:

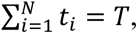

where *A_j,i_* was the abundances of the K species at grid cell *i*, respectively, and *N* was the total number of grid cells. The *S_j_* were scaling factors for each species, which rescaled their abundances by the standard deviations from the data. *t_i_* was an indicator (0/1) of transplanting at grid cell *I*, and *T* was the target number of transplants.

The optimization procedure followed these steps:

1. Begin with all available grid cells classified as “Bare”.
2. For each iteration: (a) Randomly select a batch of candidate cells (N = 20) from remaining bare cells; (b) For each candidate cell, calculate the marginal benefit of restoration by: i. Creating a temporary grid with that cell restored, ii. Recalculating Area10m for all cells within 10m radius, iii. Predicting species abundance using fitted GAMs, and iv. Computing the increase in total scaled abundance; (c) Select the cell with the highest marginal benefit for restoration, and (d) Update the grid and Area10m values.
3. Repeat until all cells are restored.

To avoid local optima, we incorporated occasional random jumps (every 50 iterations) to explore different regions of the solution space. To map the optimal restoration configurations, we extracted the grid state at the point of maximum predicted abundance and at the point where 10% of the available seascape was restored. We visualized these optimal configurations using the ’tmap’ package in R, displaying the spatial arrangement of bare, natural seagrass, and restored seagrass sites. All analyses were conducted using R version 4.4.0. We used Claude 3.7 Sonnet (Anthropic), a large language model, to assist with code development and debugging. The AI assistant helped generate the initial structure of the spatial scenarios and the optimization algorithm, which we refined and validated to ensure accurate implementation.

## Results

### The influence of habitat and seascape characteristics on animal communities and species abundance

A typical faunal community for the region, including fish, crustaceans and gastropods, were captured (Appendix S1: Table S3; Sievers et al. 2025). Species richness and Shannon diversity were highest at sites closest to mussel reefs, whilst richness was lowest within bare and 1-year-old restored seagrass sites (Appendix S1: Figure S1A, B); the opposite was observed for Evenness (Appendix S1: Figure S1C; see Appendix S1: Figures S3-S11 for plots of model fit). Evaluating beta-diversity highlighted a distinct shift in communities between those in bare and 1-year-old sites, and those in 3- and 4-year-old and natural sites (Figure 3D; Appendix S1: Table S4). These more mature seagrass sites contained more seagrass within a 10m radius and had a lower perimeter to area ratio (i.e., lower fragmentation; Appendix S1: Figure S1). Individual species responded in complex ways to restored seagrass age and seascape characteristics, and hereafter we focus on six common species. Lesser pipefish (*Sygnathus rostellatus*) abundance was highest within seagrass patches older than 1 year that had little to moderate seagrass in the surrounding 10m radius (Appendix S1: Figure S2A). Flatfish (combined *Pleuronectes platessa* and *Platichthys flesus*) abundance was highest in bare sites and at intermediate distances from mussel reefs (Appendix S1: Figure S2B).

**Figure 3.**
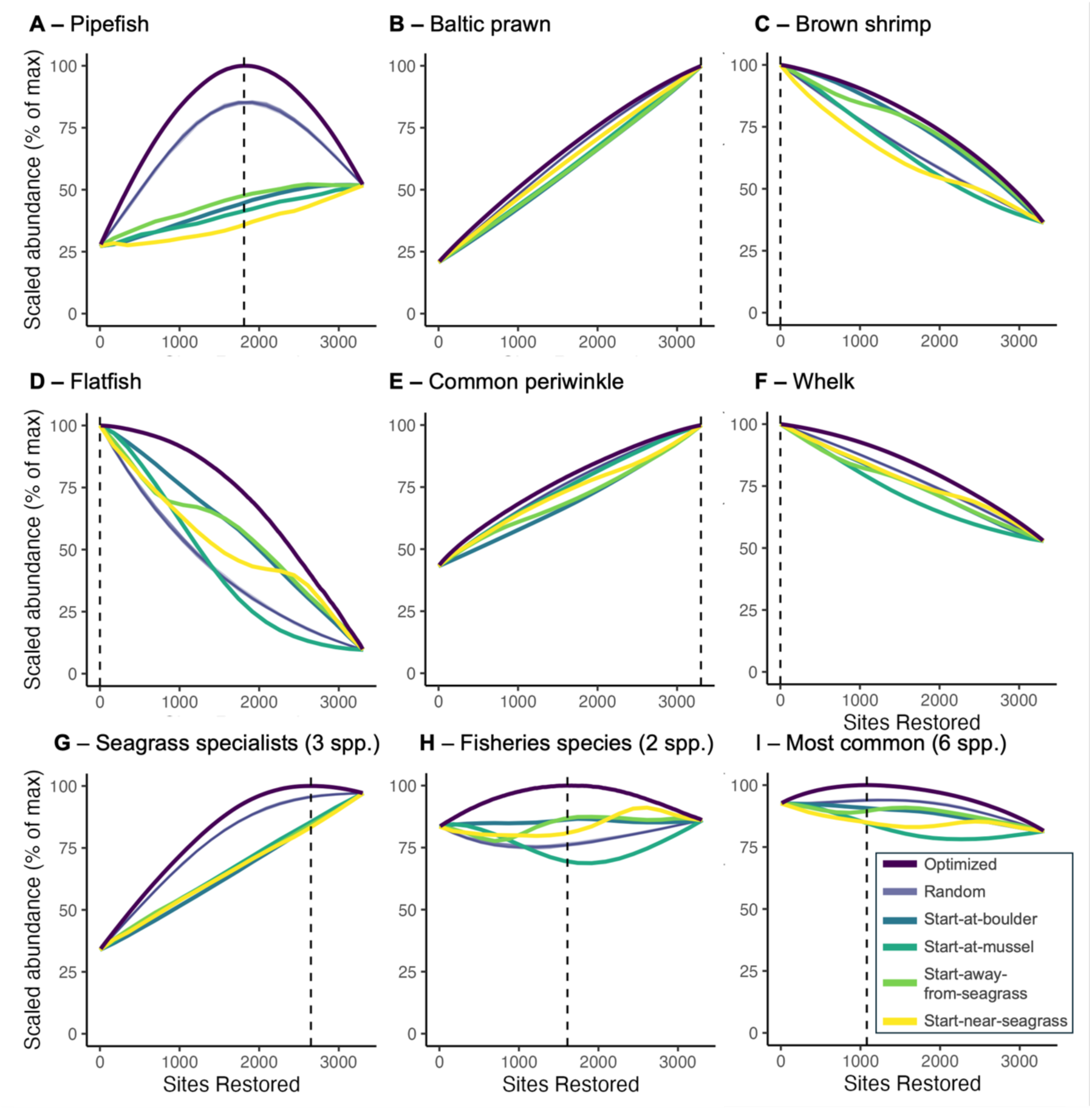
Model predictions of individual species abundances across the restoration scenarios and optimal arrangement. Scaled population abundance as a percentage of maximum for: (A) lesser pipefish (*Sygnathus rostellatus*), (B) Baltic prawns (*Palaemon adspersus*), (C) brown shrimp (*Crangon crangon*), (D) flatfish (*Pleuronectes platessa* and *Platichthys flesus*), (E) common periwinkle (*Littorina littorea*), (F) whelks (*Tritia reticulata*), (G) seagrass specialists (lesser pipefish, common periwinkle, Baltic prawns), (H) fisheries species (flatfish, Baltic prawns), and (I) most common species (all six species). Dashed lines indicate the number of sites to be restored to maximise abundance. See Appendix S1: Figure S14 for population abundance plots with marginal confidence intervals. See Appendix S1: Figure S15 for differential abundance plots for the non-optimized scenarios.

Common periwinkle (*Littorina littorea*) abundance was highest in older restored and natural seagrass sites and at sites closest to the boulder reef (Appendix S1: Figure S2C). Brown shrimp (*Crangon crangon*) abundance was highest in bare and newly restored sites, at sites farthest from the boulder reef, at sites closest to the mussel reef, and sites centrally located within the study area (Appendix S1: Figure S2D). Baltic prawns (*Palaemon adspersus*) were most abundant at sites with intermediate levels of seagrass perimeter to area ratio (Appendix S1: Figure S2E). Finally, whelks (*Tritia reticulata*) were most abundant at sites with the least amount of seagrass cover within a 10m radius and those in the northeast of the study area (Appendix S1: Figure S2F).

### Predicting animal abundance across restoration scenarios and optimizations

The optimized method unsurprisingly resulting in the maximum abundance across all objectives (Figure 3). Pipefish abundance increased as more seagrass was restored for the four scenarios that did not involve patchy transplantation, whereas for the random scenario and optimization (that involved patchy transplanting), abundance increased early and peaked at approximately half the sites being restored, then decreased (Figure 3A). As the area of restored seagrass increased, the abundance of Baltic prawns, periwinkles, and seagrass specialists increased, while the abundance of brown shrimp, flatfish and whelks decreased (Figure 3). Abundances for fisheries species and all six species objectives were hyperbolic for the optimization; maximized at an intermediate level of restoration (Figure 3H, I).

Given the optimization was always best, we describe the outcomes below as the key differences among the five remaining scenarios. For Baltic prawns, brown shrimp, periwinkles and whelks, optimal arrangements only performed slightly better than the other scenarios (Figure 3). Brown shrimp were most abundant for the scenarios start- near-boulder-reef and start-away-from-seagrass (Figure 3C). The steep decline in flatfish abundance observed during the intermediate stages of the start-near-mussel-reef scenario (Figure 3D) can be explained by the unimodal relationship between abundance and distance to mussel reef (Appendix S1: Figure S2D), whereby sites with the highest predicted abundance undergo a transition from bare to seagrass during the intermediate steps for this scenario, resulting in a rapid loss of sites with the most flatfish. Whelk abundance was slightly lower for the start-near-mussel-reef scenario (Figure 3F), partly driven by the conversion of bare sites in the northeast where the species was most abundant (Appendix S1: Figure S2F). Seagrass specialists and the six most common species objectives both had highest abundances for the random scenario (Figure 3G, I), whilst fisheries species abundance was maximized across different scenarios (not including random) depending on how much of the seascape was restored (Figure 3H). For fisheries species and the six most common species, the start-at-mussel scenario generally performed the worst (Figure 3H, I).

### Optimal transplant arrangements

We first map the optimal arrangement of seagrass transplants based on restoring 10% of the seascape. All nine objectives showed that some level of patchy planting was best, with the locations of these planted areas differing substantially across objectives (Figure 4). Restored sites for Baltic prawns, flatfish and whelk were generally far away from mussel and boulder reefs, whilst brown shrimp and periwinkles were near (Figure 4).

**Figure 4.**
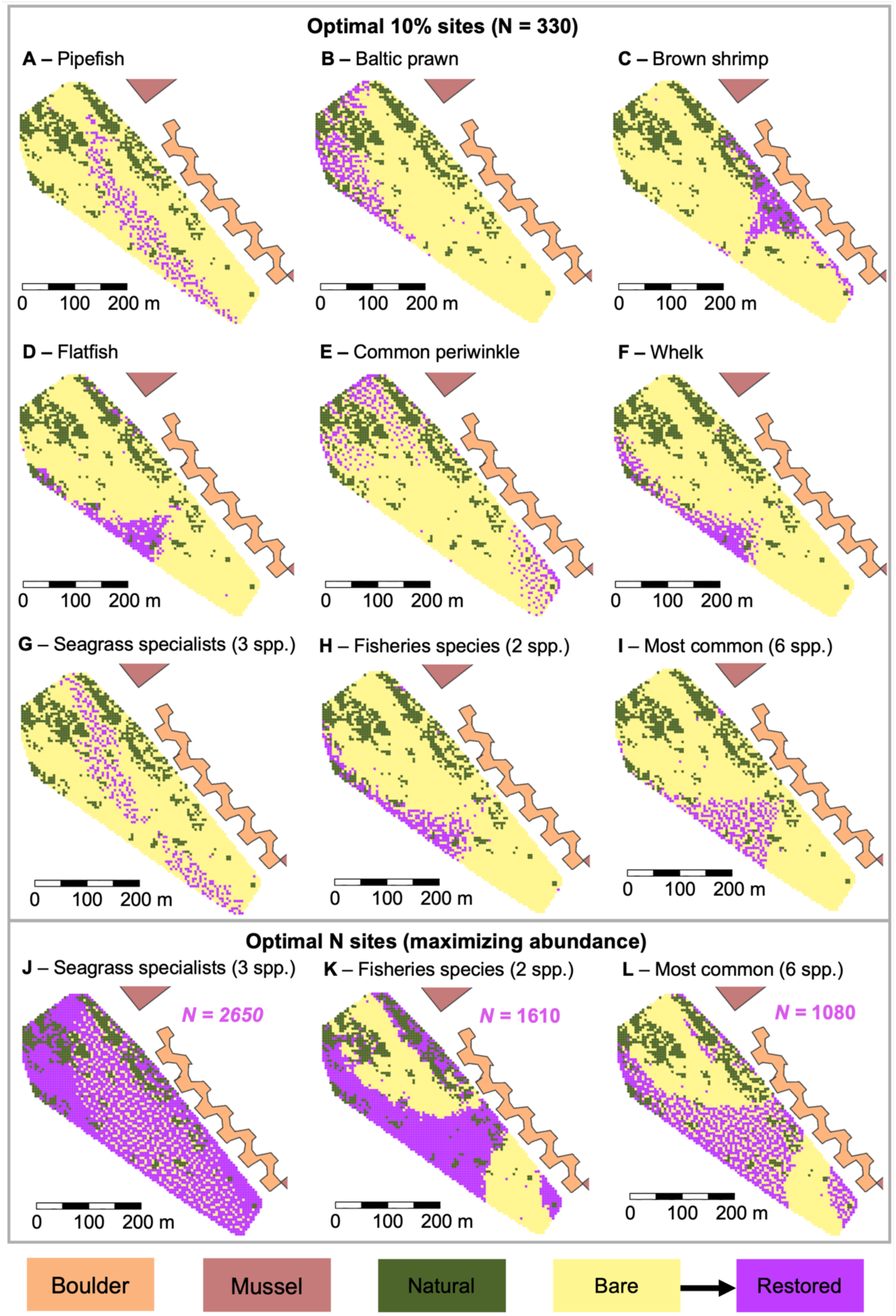
Optimizing restoration site selection for alternate objectives. Optimal position of sites to maximise the objective when restoring 10% of the available sites (A– I), and when no restriction is placed on how many sites can be restored (i.e., maximising abundance; J –L). The boulder reef (orange) and mussel reefs (salmon) are also shown. See Appendix S1: Figure S16 for marginal model predictions used to calculate the marginal value of restoration used in the optimization for the three multi-species objectives. For maps of maximising abundance pipefish, see Appendix S1: Figure S13; other species abundances were maximised at either all or none of the seascape. Lesser pipefish (*Sygnathus rostellatus*); Baltic prawns (*Palaemon adspersus*); brown shrimp (*Crangon crangon*); flatfish (*Pleuronectes platessa* and *Platichthys flesus*); common periwinkle (*Littorina littorea*); whelks (*Tritia reticulata*); seagrass specialists (lesser pipefish, common periwinkle, Baltic prawns); fisheries species (flatfish, Baltic prawns), and; the most common species (all six species).

Sites for pipefish and seagrass specialists were at intermediate distances from these other restored habitats, whilst sites for fisheries species and the most common species objectives were farther away (Figure 4). We found minimal spatial overlap in the optimal 10% of sites for some objectives; for example, only 37 out of the possible 330 restored sites overlapped between the fisheries species and pipefish objectives (Appendix S1: Figure S12). Maps of the optimal arrangement to maximise abundances (i.e., no restriction on the amount of the seascape restored; Figure 4J–L) for most species where either all sites restored (Baltic prawns and periwinkles) or no sites restored (brown shrimp, flatfish and whelks), which aligns with abundance plots (Figure 3). Pipefish abundance was maximized at 55% of the sites being restored (1810 out of 3310 sites restored) and was highly patchy (Appendix S1: Figure S13), whilst seagrass specialists were maximized at 80%, fisheries species at 49%, and the most common species at 33% (Figure 4). Fisheries species and the most common species objectives were maximized by restoring very similar areas of the seascape – at areas either far or near the mussel reefs – but the latter involved a patchy planting arrangement (Figure 4).

## Discussion

We show how empirical data from animal monitoring can be used to develop spatially explicit predictions of ecological outcomes for different seagrass planting arrangements, including an optimal design. Clear and diverse relationships between abundances and seascape characteristics in this multi-habitat restoration seascape provided a strong basis for showing how strategic seagrass restoration can optimize outcomes for biodiversity and target species through spatially strategic planting of seagrass. Outcomes can vary greatly depending on where restoration occurs (even at small spatial scales), and thus, optimal planting arrangements were discernably different across objectives, highlighting the importance of restoration designs and setting clear objectives. Despite this variability, a patchy planting arrangement was a well-supported strategy for most objectives.

Quantifying species abundances was integral to developing models to predict outcomes from various planting arrangements and the spatial optimization outputs. The distinct shift in community composition between bare/newly restored sites and mature seagrass beds aligns with previous research showing that faunal colonization of restored habitats can occur rapidly (Lefcheck et al. 2017, Beheshti et al. 2021, Tanner et al. 2021, Steinfurth et al. 2022, Gagnon et al. 2023, Sievers et al. 2025). Proximity to other restored habitats – mussel and boulder reefs – also influenced community metrics, with richness and Shannon diversity highest near mussel reefs. Individual species also showed markedly different responses to restoration and seascape characteristics, reflecting diverse habitat requirements and resource needs. Lesser pipefish, for example, were most abundant in established seagrass beds (natural and restored), but showed reduced abundance in areas with high seagrass coverage within a 10m radius, possibly indicating a preference for habitat edges or meadows of moderate density that balance preferences for feeding efficiency and predator avoidance (Flynn and Ritz 1999, Sievers et al. 2025). In contrast, flatfish – an important fisheries species – were most abundant in bare sites and at intermediate distances from mussel reefs. Collectively, this suggests that alternate restored habitats may provide additional and/or complementary resources that benefits some species and can enhance local biodiversity, and is consistent with work showing that habitat connectivity within coastal ecosystems plays a key role in supporting some species and enhancing positive facilitative interactions (Vozzo et al. 2023, Silliman et al. 2024).

The distinct species-specific responses resulted in discernibly different predicted outcomes across our restoration scenarios. The optimization analyses demonstrated that strategic, patchy planting configurations may offer more beneficial outcomes for animal populations compared to uniform, continuous seagrass coverage. This aligns with current seagrass restoration strategies that recognize the benefits of patchy planting (e.g., reduced restoration costs and workload, as it leverages the natural growth capabilities of seagrass; Flindt et al. 2024), and aligns with how natural seagrass may recruit from seed into bare areas in many regions (Olesen and Sand-Jensen 1994, Orth et al. 1994). This finding also echoes work that shows that small, fragmented meadows can support equivalent biodiversity and functional diversity to larger, less fragmented meadows (Lefcheck et al. 2016, Gagnon et al. 2023). While it is anticipated that seagrass will naturally grow and fill in these gaps, potentially affecting our predictions for some objectives, based on the fragmentation of natural patches in many regions (including Vejle fjord), it is also possible that the meadow will continue to have some degree of fragmentation due to hydrodynamics and storms (Frederiksen et al. 2004). Our results also point towards the plausible effects that seascape homogenization has on animals, whereby fully converting bare areas to any one habitat type is not necessarily ideal and a heterogeneous seascape may benefit animal communities. This matches work at larger spatial scales, for instance with mangroves, where estuaries with large mangrove extents tend to have very homogeneous seascapes, leading to lower fish abundance and diversity (Henderson et al. 2020). This finding also relates to area-based restoration targets, whereby larger areas could be achieved that produce better outcomes for fewer resources. Ultimately, our results emphasize the complexity of designing restoration projects that aim to benefit multiple species simultaneously, the importance of clear restoration objectives when planning habitat/planting configurations, and the inherent tradeoff that exists between optimizing animal outcomes with processes enhancing restoration success.

### Limitations and future directions

Our analyses and approach are based on a single multi-habitat restoration seascape. However, our intention was not to provide specific guidance for seagrass restoration in Vejle Fjord, but rather to showcase how animal monitoring data can be leveraged to develop spatially explicit predictions and optimise objective-dependent restoration planning. Further, while we focused on fine-scale habitat interactions and effects (i.e., 10 m radius, and restored plot size of ∼20 m^2^), many seascape studies demonstrate both positive and negative influences of seascape characteristics at scales of 10s to 1000s of metres (Vozzo et al. 2024). Future studies would thus benefit from collecting temporal data across multiple locations with different combinations and configurations of habitat types, and incorporate data at spatiotemporal scales most relevant to the study system and restoration objectives. This would provide greater capacity to test the benefits of multi-habitat restoration approaches and allow for more robust predictions of animal responses.

Our models could also be improved by incorporating additional parameters where data exists, such as on environmental factors (e.g., wave exposure, water quality), economic and logistical considerations, and habitat suitability of the habitat forming species. For instance, analyses could explore how optimal configurations change when accounting for plant mortality, where seagrass grows best, where planting is cheapest, or where water or sediment quality thresholds are not breached (Thom et al. 2018, Flindt et al. 2024).

Scenarios could also examine the implications of adding or removing one or more habitat types (e.g,. mussel and boulder reefs), which would help identify the most effective combinations of habitats for achieving specific restoration objectives while maximising cost-effectiveness for multi-habitat projects. Notably, for this type of modelling, manipulative and/or fully orthogonal tests of multi-habitat restoration are preferred before definitive conclusions can be drawn; this may be currently outside the realm of what is possible given the lack of multi-habitat seascape restoration (Olds et al. 2016, Vozzo et al. 2023).

One of the multi-species objectives we focus on is based on the standardized abundance of the six most common species, a group consisting of the three seagrass specialists and three species that are more abundant in bare sand. It is, however, highly probable that a greater proportion of the species surveyed (but not modelled here due to data paucity) prefer seagrass and would directly benefit from its restoration. Indeed, species richness and diversity relationships here and in previous work (Sievers et al. 2024) confirm this, suggesting that the community-wide biodiversity benefits of restoring seagrass are likely even greater than predicted here. Furthermore, although we see immediate decreases in the predicted abundance of some species (e.g., flatfish, brown shrimp) as seagrass is restored, we might not expect this in reality. Currently, these models and predictions do not account for ‘overflow effects’ or subsidies provided by the addition of more seagrass habitat; instead, they are based on the distribution of animals in the seascape at the time of monitoring, and thus reflect habitat occupation. Our models may thus underestimate population increases following seagrass restoration, for instance, if the carrying capacity of the whole system increases more than predicted due to these extra subsidies.

Conversely, the models do not impose limits on carrying capacity or lagged density effects (e.g., due to competition or disease) so may also overestimate true abundances. More sophisticated models that account for species-habitat interactions, carrying capacity, trophic subsidies, life-history-dependent habitat preferences, and species interactions are becoming easier to implement and, with sufficient data, may produce more accurate predictions to inform restoration decisions and action (Heymans et al. 2016, Clark et al. 2017).

### Applying our approach in practice

Our framework for informing multi-habitat restoration, while demonstrated here using data from an established multi-habitat restoration project, can be adapted for use in new restoration initiatives through strategic data collection. For multi-habitat projects especially, a key challenge for practitioners will be obtaining relevant data to inform spatial planning, particularly given the current scarcity of multi-habitat restoration sites (Vozzo et al. 2023) and the general lack of animal data from restored sites (Sievers et al. 2024). However, this limitation can be addressed by increasing animal monitoring efforts and carefully selecting sites that span different natural configurations of target habitats (Gilby et al. 2019, Henderson et al. 2019). For instance, practitioners can leverage existing seascapes where multiple habitat types co-occur naturally at varying distances and arrangements to understand how spatial configuration influences target species.

Similar approaches have been used to quantify seascape effects on fish assemblages (Olds et al. 2016) and inform restoration planning (Gilby et al. 2021). Even single-habitat restoration projects that are cognizant of surrounding habitat types can provide valuable insights into species-habitat relationships at relevant scales (Gilby et al. 2018, Vozzo et al. 2024). To facilitate the adoption of our approach, we provide our code that can be adapted for different locations, contexts and species assemblages. While the specific optimal configurations will undoubtedly vary among locations and target species, our statistical framework offers a structured way to incorporate empirical data into spatial planning, regardless of whether that data originates from existing restoration projects, natural reference sites, or a combination of both.

As restoration efforts continue to expand globally in response to international commitments, evidence-based approaches will become increasingly valuable for maximizing the return on restoration investments and achieving project objectives (Duarte et al. 2020, Elliott et al. 2022). Our study presents a proof-of-concept framework for incorporating animal monitoring data into spatial planning for coastal restoration projects. The utility of predictive models lies in their ability to inform strategies for spatial prioritization, which can then be translated into practice as simple guidelines. Our optimization boils down to the recommendations of ‘plant patchy’ and, for some species such as brown shrimp and periwinkles, ‘plant for connectivity of the target species’, whereby these species showed benefits of patchy planting near alternate restored habitats. Importantly, not all species were most abundant near alternate restored habitats, including fished species, so defining objectives is of the utmost importance. The framework presented here can be adapted to different ecosystems, sites and objectives, providing a valuable tool for practitioners working to enhance biodiversity and ecosystem services through habitat restoration. Ultimately, by combining data on species-habitat relationships with the spatial arrangement of restored habitats, these predictive tools can help practitioners optimize restoration strategies, which along with financial, cultural and logistical considerations, can help achieve project-specific objectives such as enhancing biodiversity or fisheries production.

## Acknowledgments

MS was supported by a funded by Australian Research Council (ARC) Discovery Early Career Researcher Award no. DE220100079. CJB was supported by a Future Fellowship (FT210100792) from the Australian Research Council. BN, RCS, MRF and TLB was supported by the “Healthy Vejle Fjord” project funded by the Velux Foundations call “Et Hav i Balance” (Sund Vejle Fjord (25098)). We acknowledge the use of Claude 3.7 Sonnet (Anthropic) for assistance with developing and refining code.

## Author Contributions

MS, CJB, JAR, BN, RCS, MRF, RMC conceptualized the study; Field work was conducted by MS, JAR, BN, RS, RMC and resourced by MS, MF, RMC; Samples were processed and analyzed by MS, JAR, RCS, BN; Data analysis was led by MS and CJB with contributions from all authors; All authors interpreted results; manuscript writing was led by MS with contributions from all authors. All authors approve submission of this manuscript.

