## Supplementary Material for "Optimizing seagrass planting arrangements for animal benefits in a multi-habitat restoration seascape"

**Appendix S1**

Optimizing seagrass planting arrangements for animal benefits in a multi-habitat restoration seascape

Figures S1 to S19

Tables S1 to S5

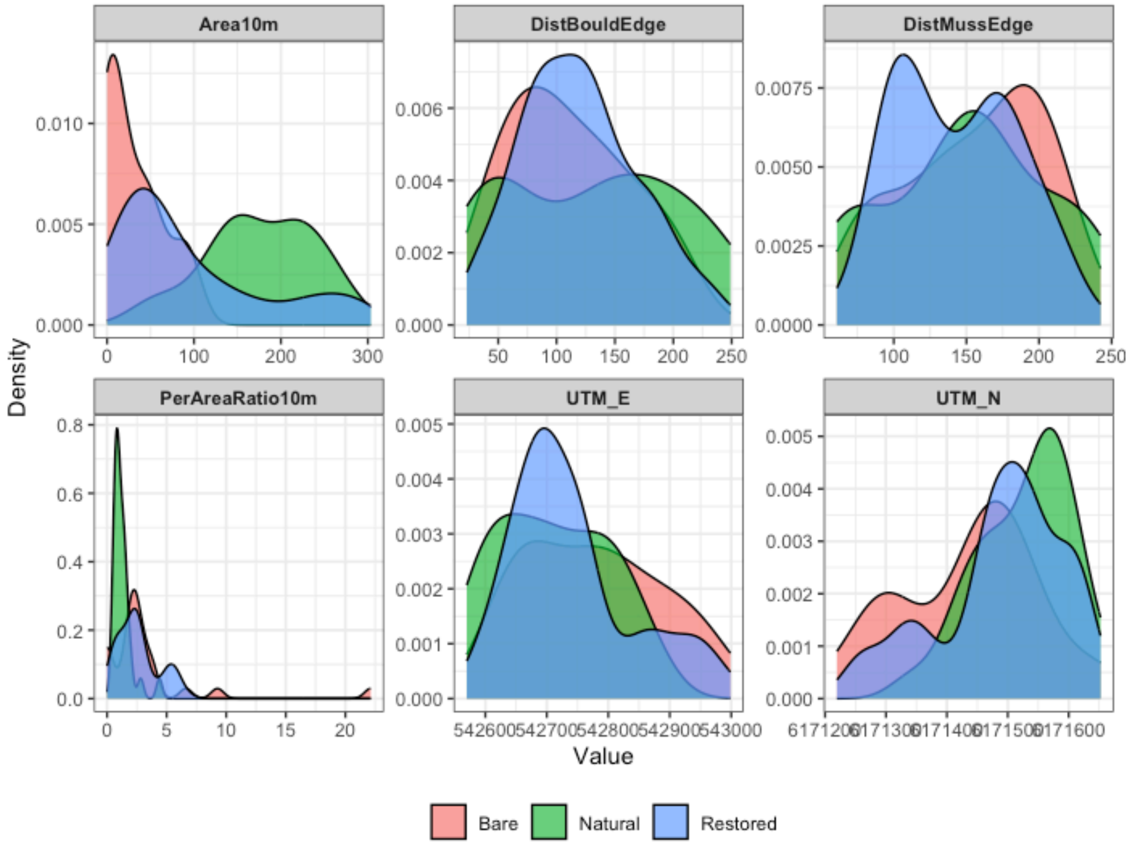


**Figure S1. Density distribution of spatial covariates across Treatment.** Distribution plots show the spread of key spatial variables across bare (pink), natural seagrass (green), and restored seagrass (blue) sites. Covariates include seagrass area within 10m radius (Area10m), distance to boulder reef edge (DistBouldEdge), distance to mussel reef edge (DistMussEdge), perimeter-to-area ratio within 10m (PerAreaRatio10m), and UTM coordinates (UTM_E and UTM_N). The substantial overlap in distributions for distance-based covariates indicates balanced spatial sampling across treatment types, supporting the validity of our observational approach.

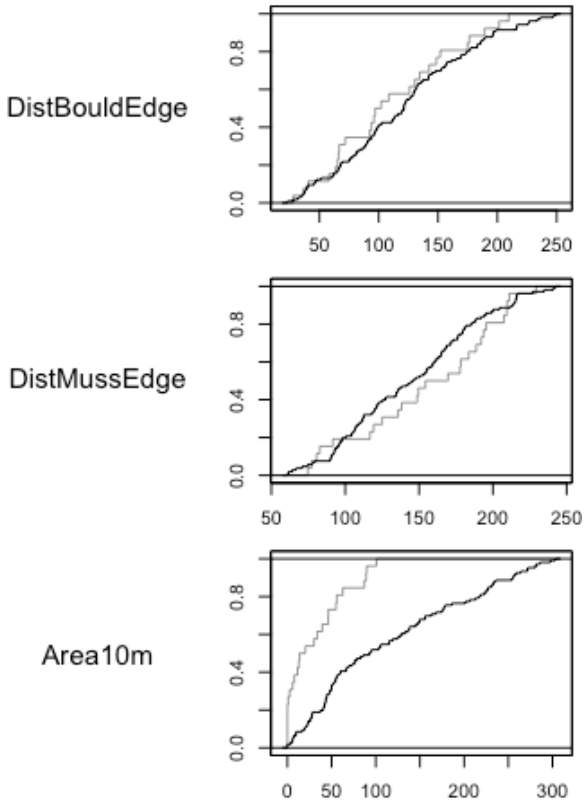


**Figure S2. Empirical cumulative distribution function (eCDF) plots for key spatial covariates used in predictive models.** Plots compare the distributions of distance to boulder reef edge (DistBouldEdge), distance to mussel reef edge (DistMussEdge), and seagrass area within 10m radius (Area10m) after matching. Black lines represent seagrass sites while gray lines represent bare sand sites. The similar trajectories of the lines demonstrate balanced spatial coverage across our sampling design, reducing potential spatial bias in our observational study.

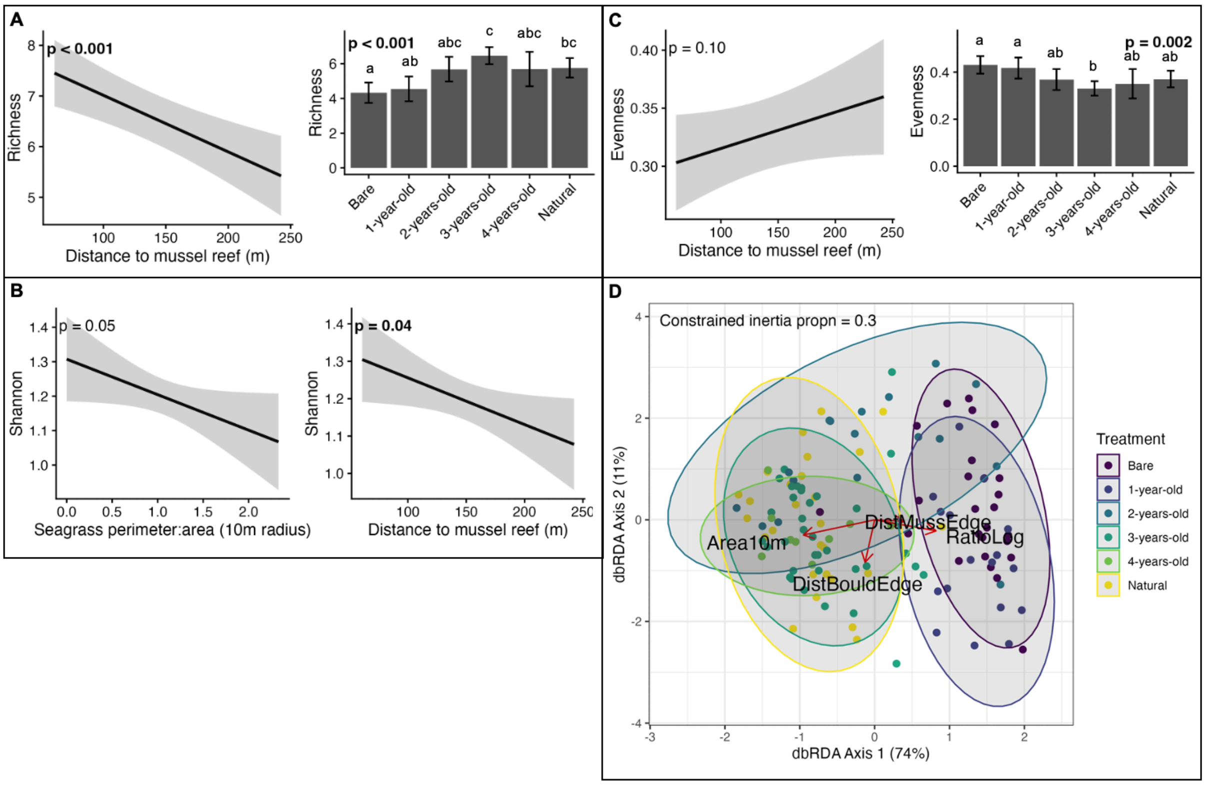


**Figure S3. Relationship between (A) species richness, (B) Shannon diversity, (C) Evenness, and (D) beta diversity and habitat/seascape characteristics.** Plots A-C are based on best-fitting generalized additive models (GAM) with the statistical significance of the factors shown. Error bars and ribbons are 95% confidence intervals. Plot D is a based on a distance-based redundancy analysis (dbRDA) for the top 10 most abundant species and a Bray-Curtis dissimilarity matrix (Area10m – area of seagrass within a 10m radius; DistBouldEdge – distance to the boulder reef; DistMussEdge - distance to the mussel reef; RatioLog – log-transformed perimeter to area ratio). See Table S5 for ANOVA outputs. See Figure S17 for beta-diversity model diagnostics.

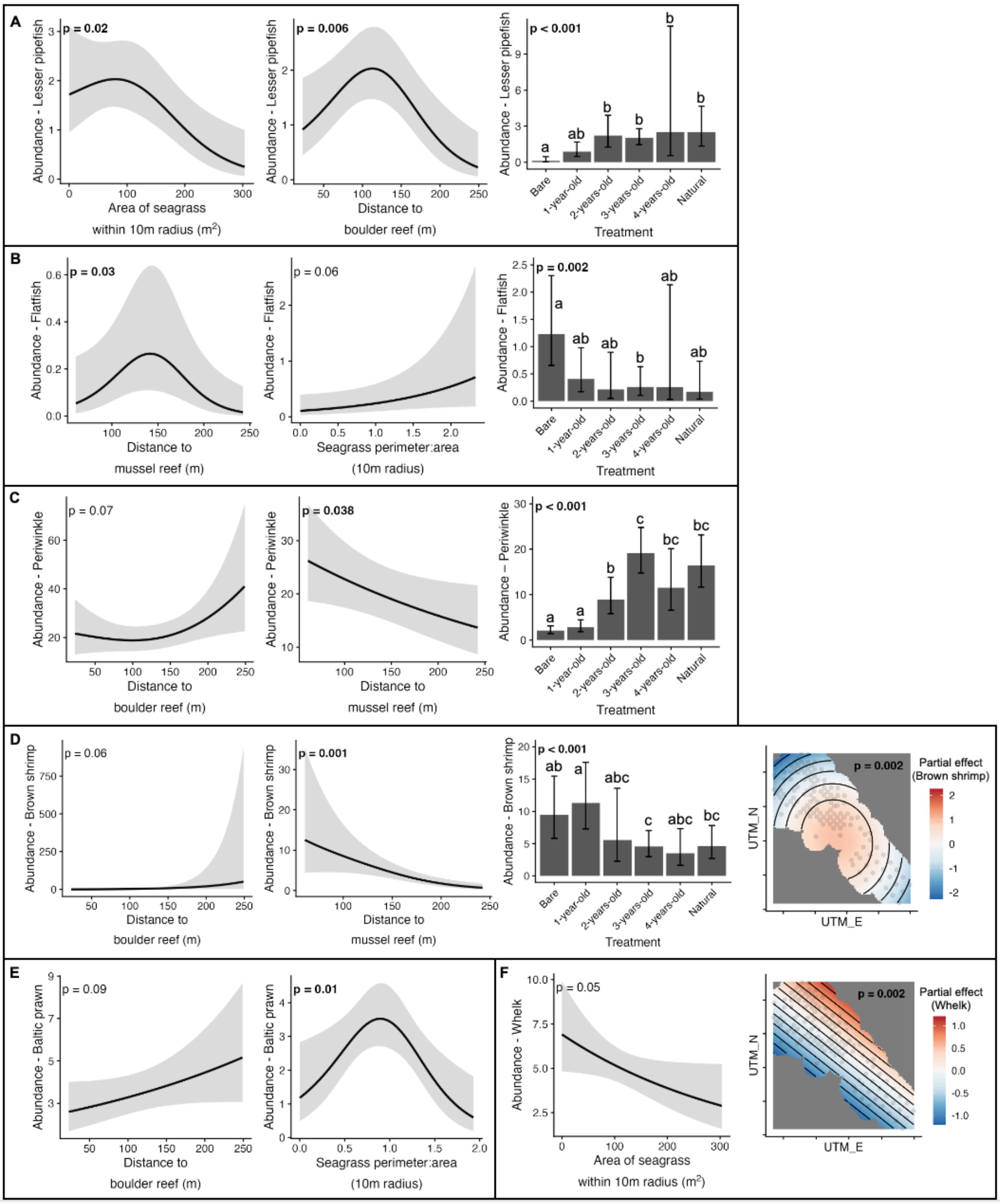


**Figure S4. Relationship between species abundances and habitat/seascape characteristics for the 6 most common species (A-F).** Plots are based on best-fitting generalized additive models (GAM) with the statistical significance of each of the remaining factors shown. Error bars and ribbons are 95% confidence intervals. For spatial random effects plots (E and F), values are partial effect sizes. See Table S5 for ANOVA outputs.

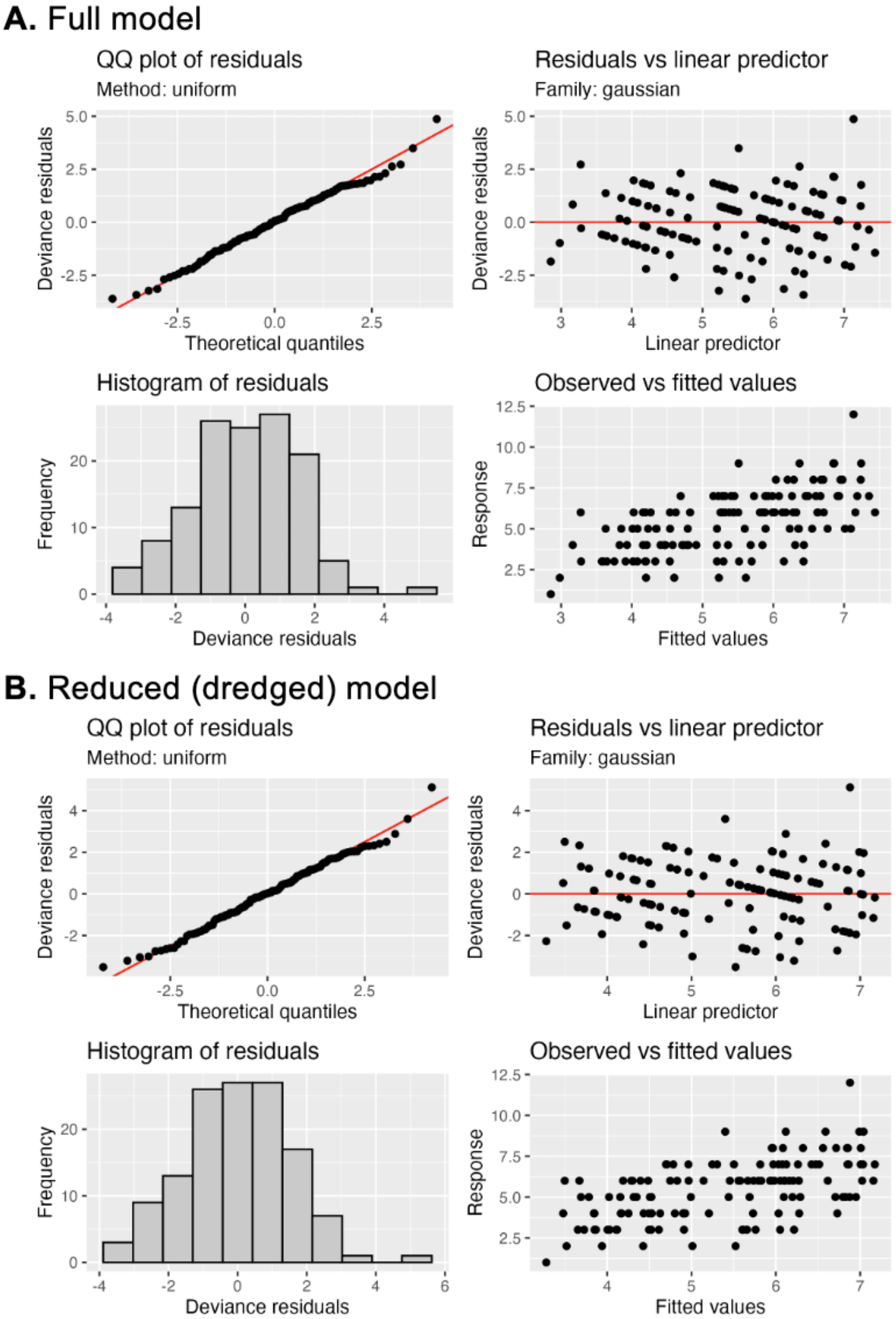


**Fig. S5.** GAM diagnostic plots for species richness, for both the full model and the best-fitting reduced (dredged) model.

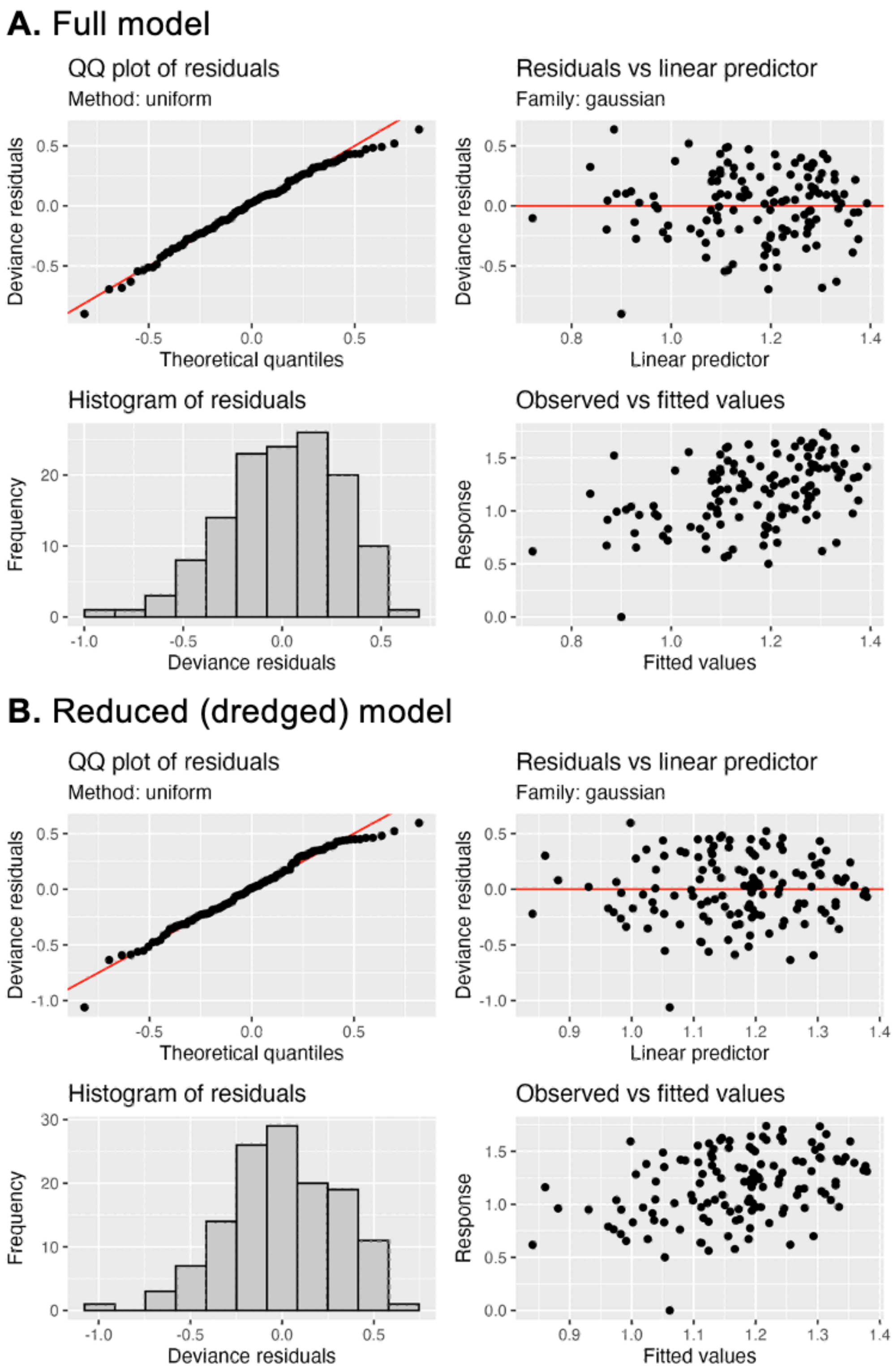


**Fig. S6.** GAM diagnostic plots for Shannon diversity, for both the full model and the best-fitting reduced (dredged) model.

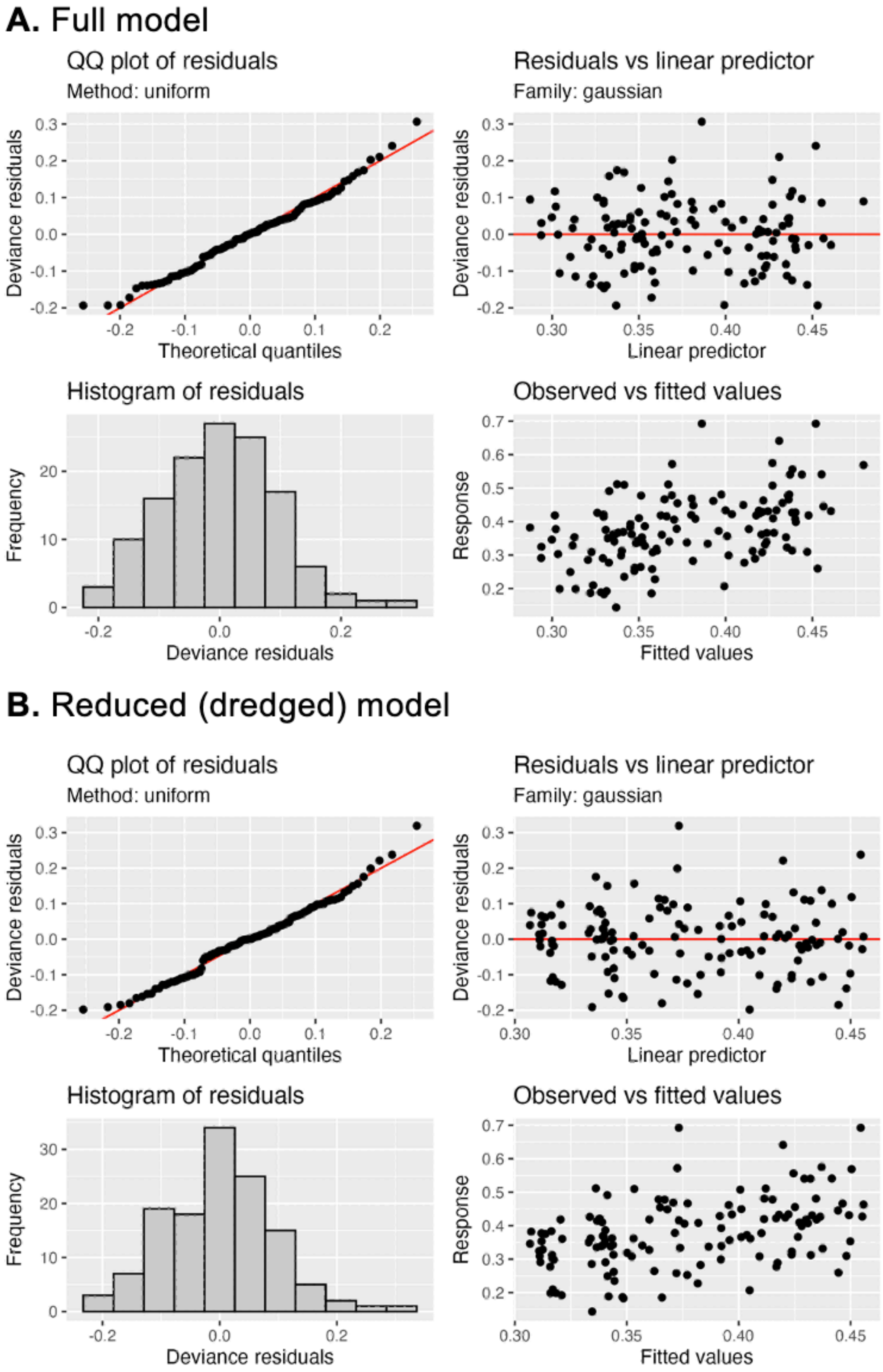


**Fig. S7.** GAM diagnostic plots for evenness, for both the full model and the best-fitting reduced (dredged) model.

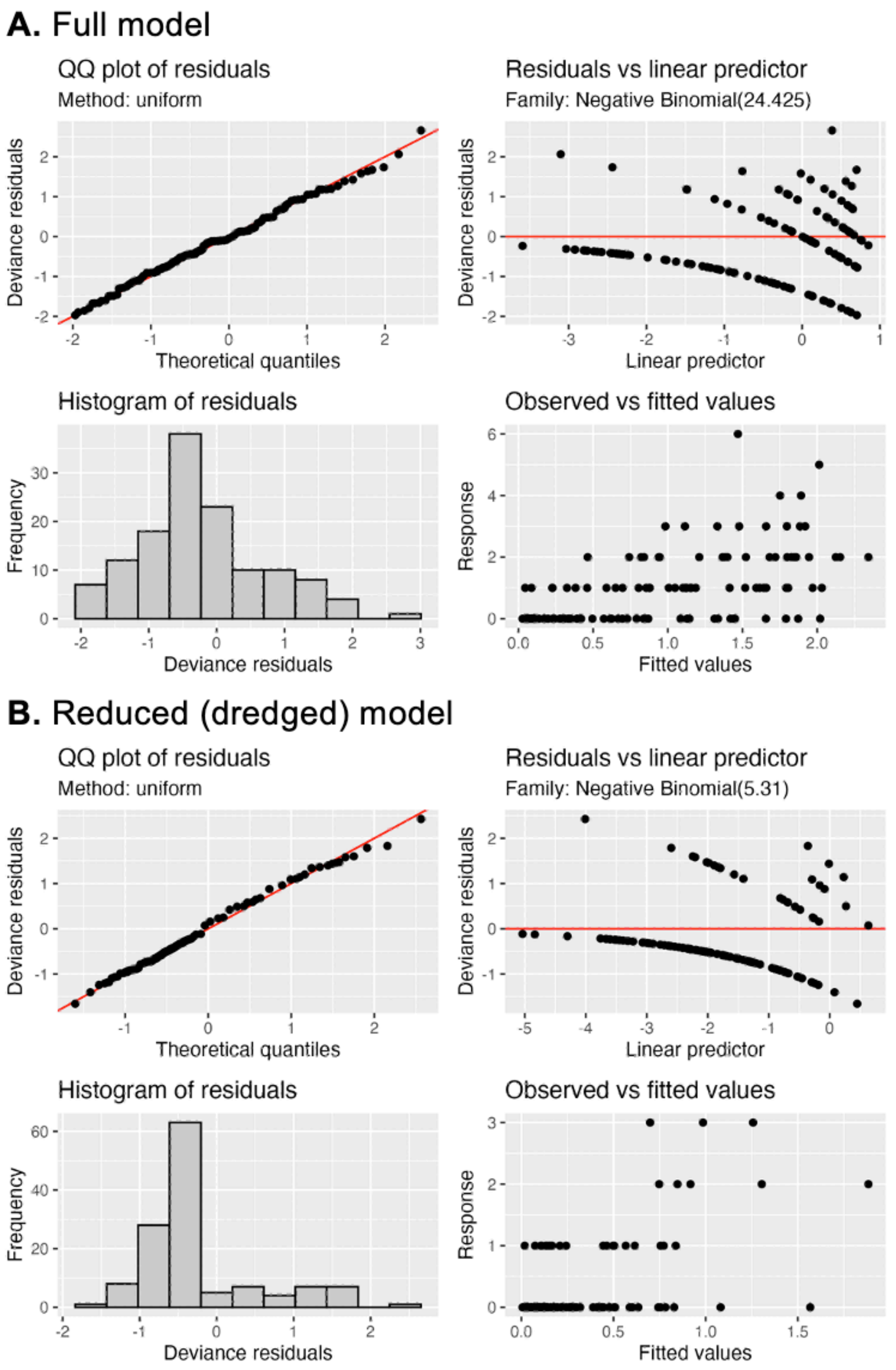


**Fig. S8.** GAM diagnostic plots for lesser pipefish abundance, for both the full model and the best-fitting reduced (dredged) model.

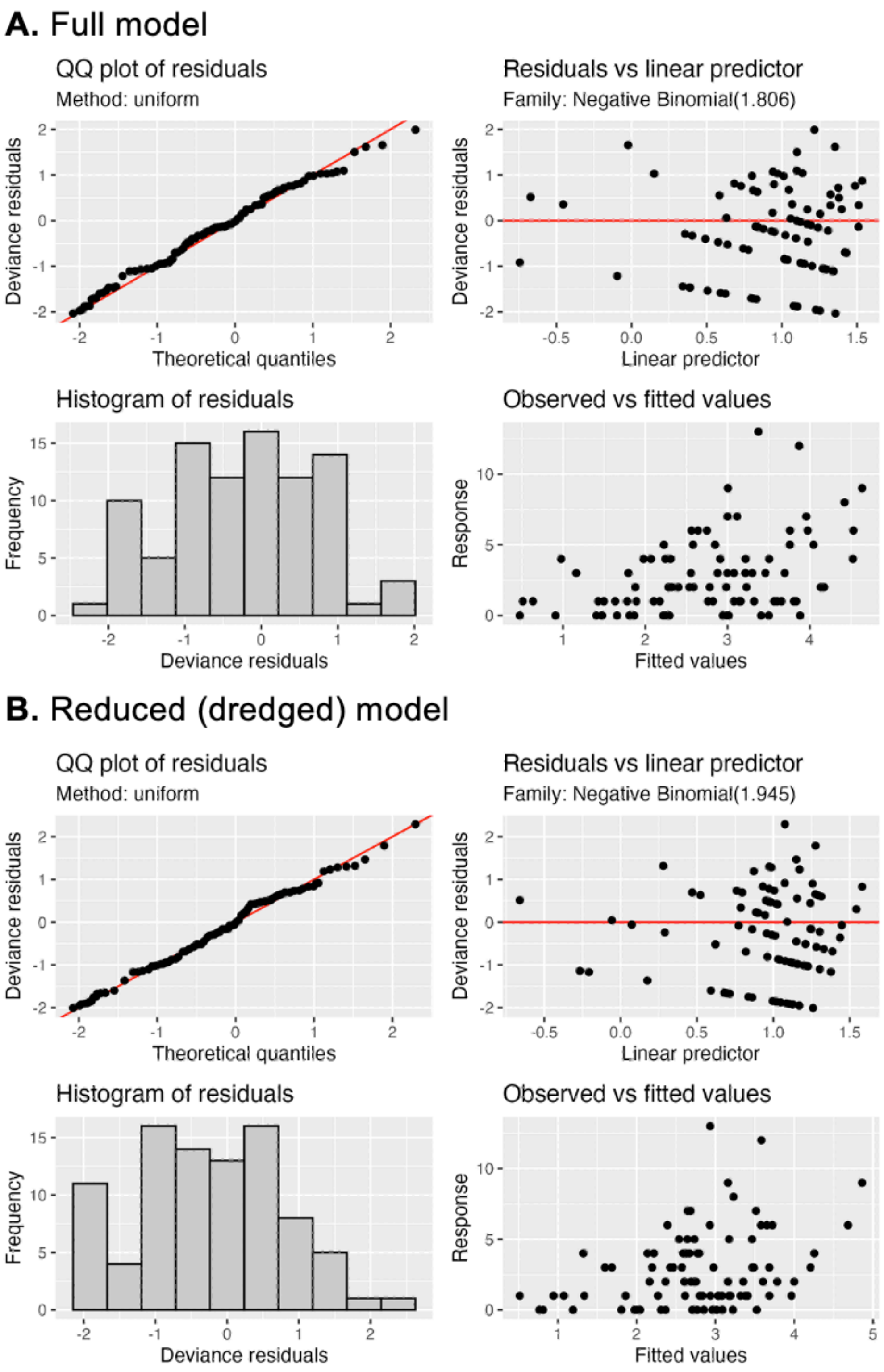


**Fig. S9.** GAM diagnostic plots for Baltic prawn abundance, for both the full model and the best-fitting reduced (dredged) model.

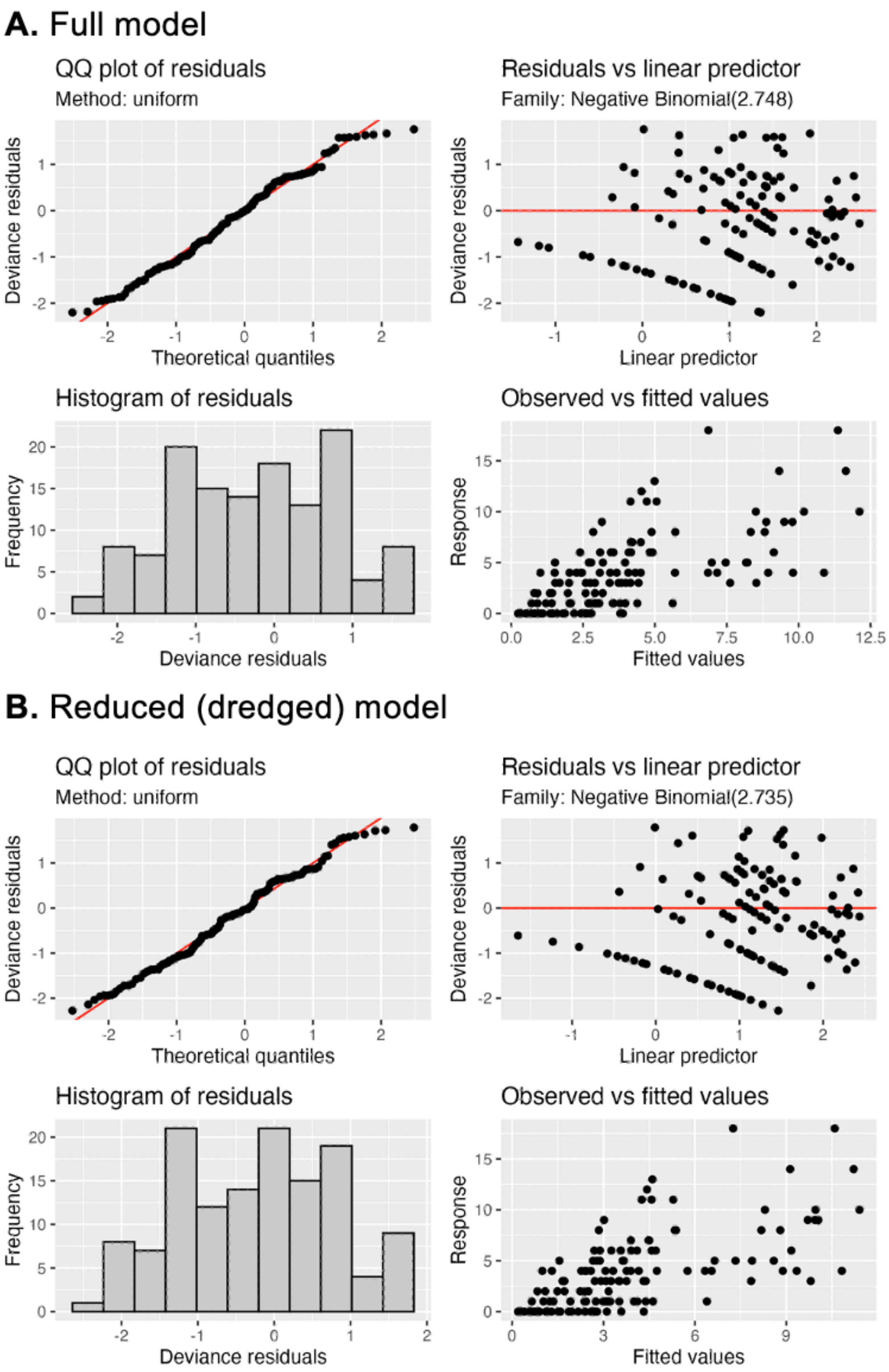


**Fig. S10.** GAM diagnostic plots for brown shrimp abundance, for both the full model and the best-fitting reduced (dredged) model.

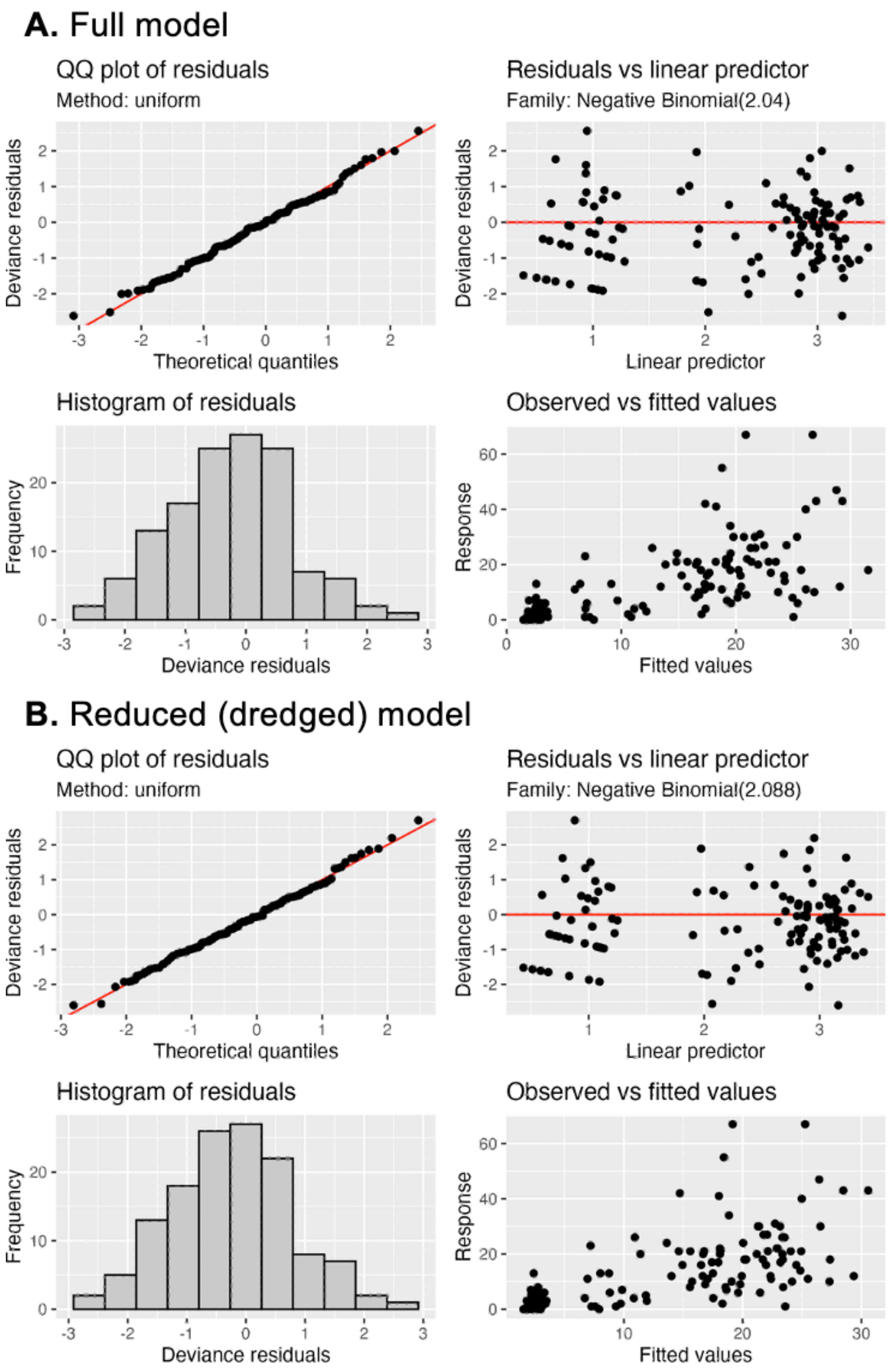


**Fig. S11.** GAM diagnostic plots for common periwinkle abundance, for both the full model and the best-fitting reduced (dredged) model.

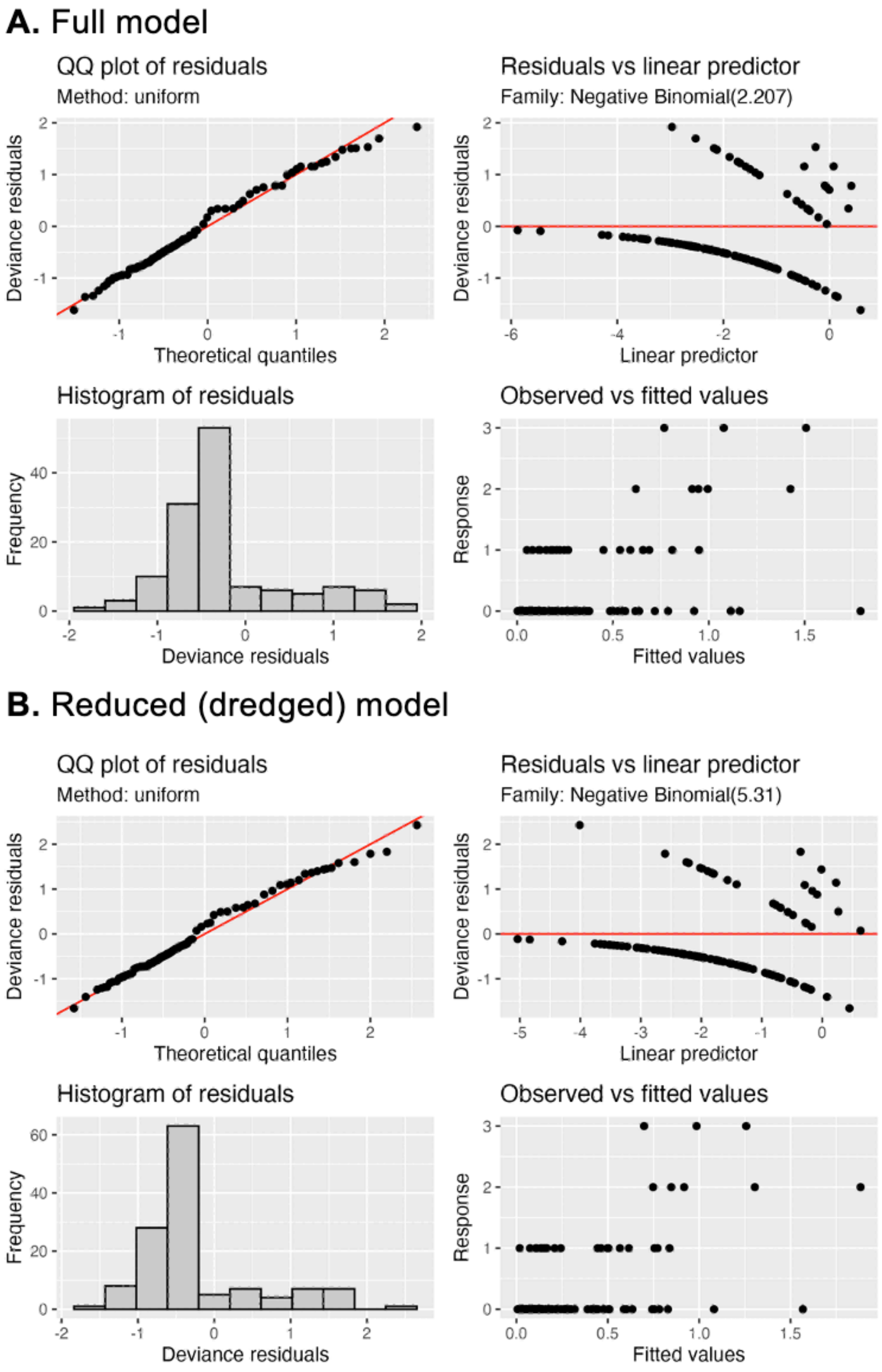


**Fig. S12.** GAM diagnostic plots for flatfish abundance, for both the full model and the best-fitting reduced (dredged) model.

**A.** Full model

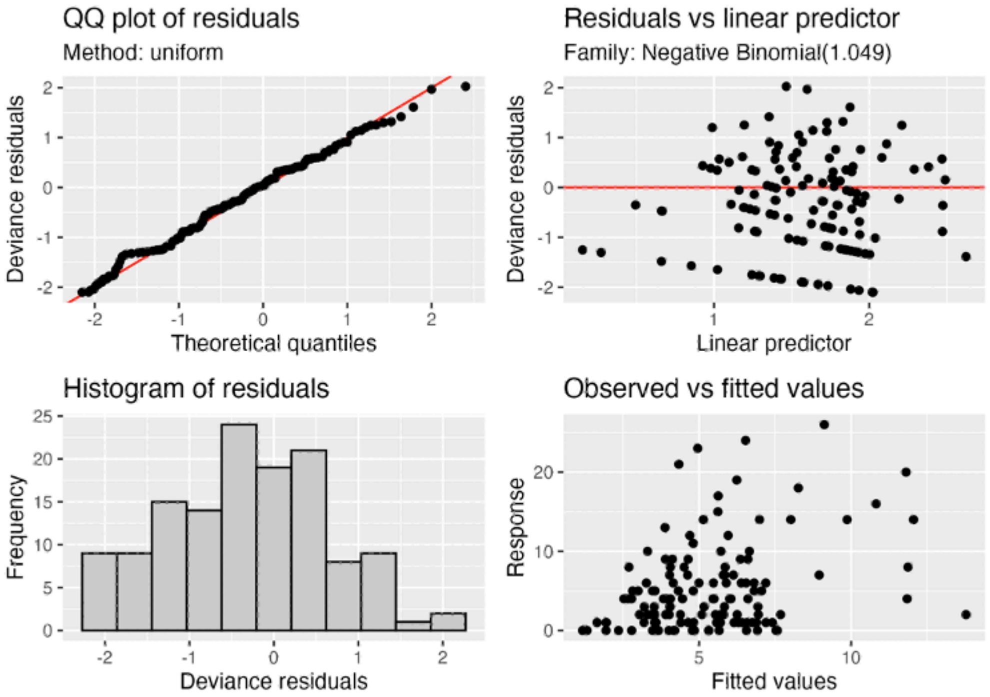


**B.** Reduced (dredged) model

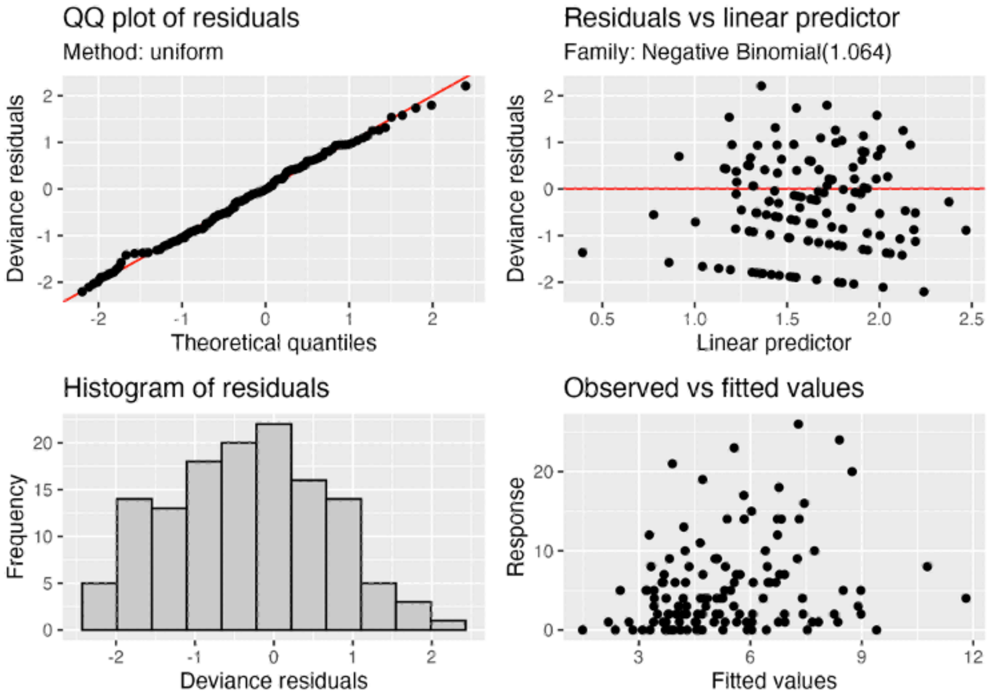

**Fig. S13.** GAM diagnostic plots for whelk abundance, for both the full model and the best-fitting reduced (dredged) model.

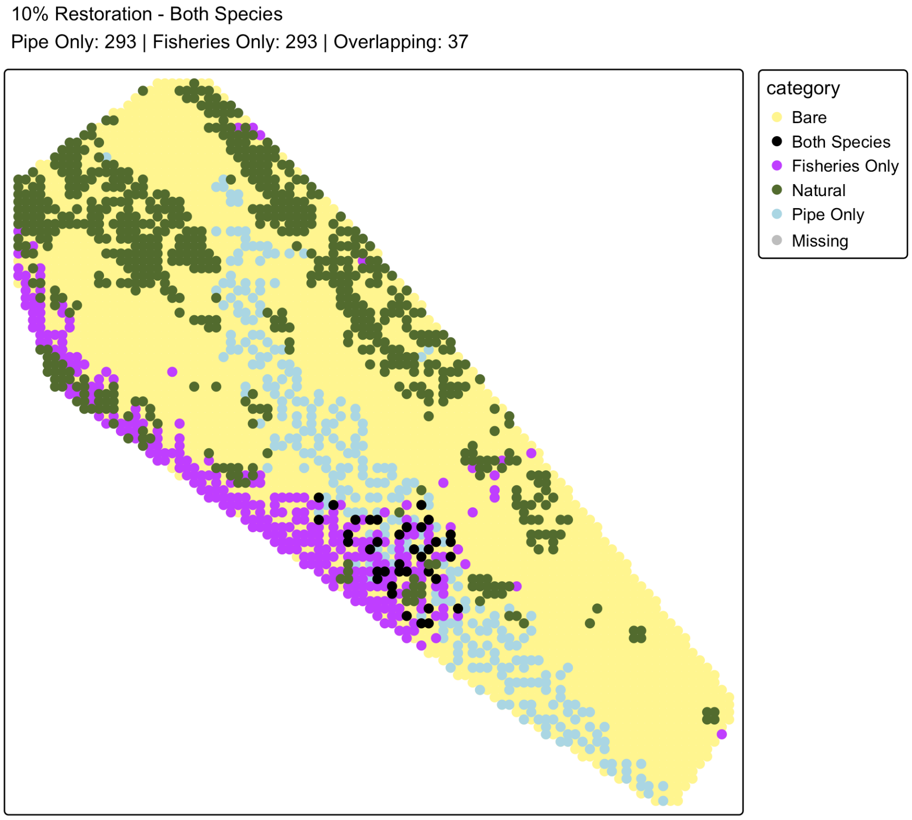


**Fig. S14.** The optimal position of sites to maximise two objectives – fisheries (flatfish, Baltic prawns) and lesser pipefish - when restoring 10% of the available sites, to identify sites that benefit both objectives.

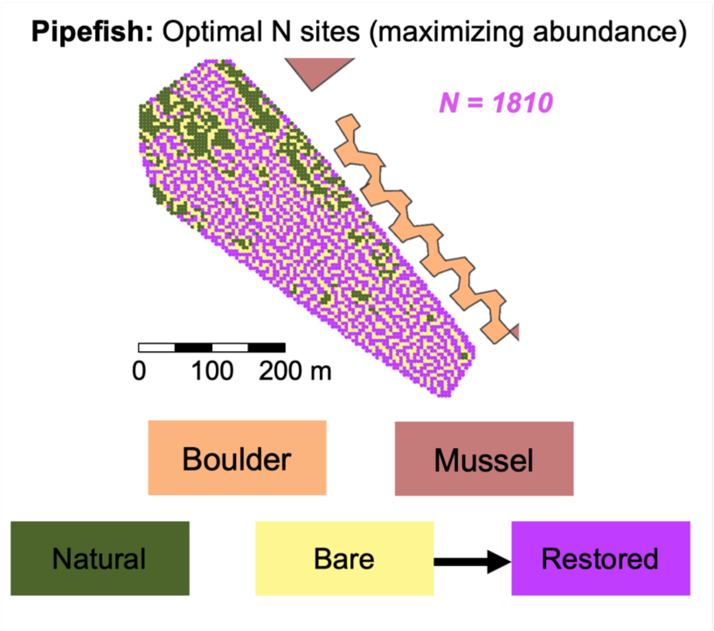


**Fig. S15. Optimizing restoration site selection for lesser pipefish (*Sygnathus rostellatus*).** Optimal position of sites to maximise the objective when when no restriction is placed on how many sites can be restored (i.e., maximising abundance). The boulder reef (orange) and mussel reefs (salmon) are also shown.

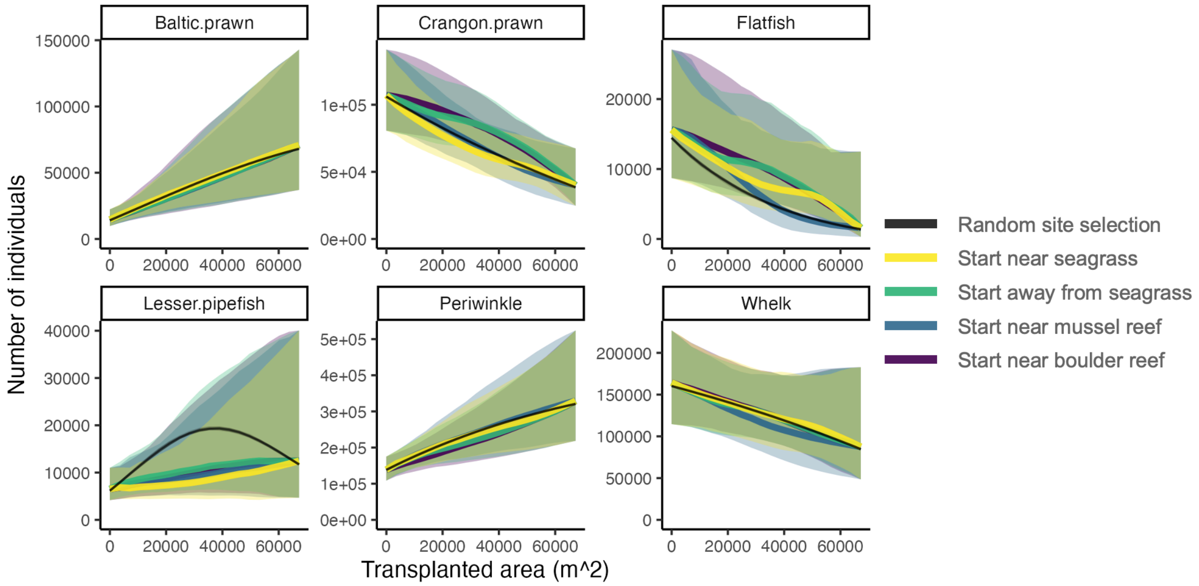


**Fig. S16. Model predictions of population abundance for the six most common species**, relative to the estimated abundance within the seacape without any restored seagrass, for five restoration scenarios with 95% marginal confidence intervals.

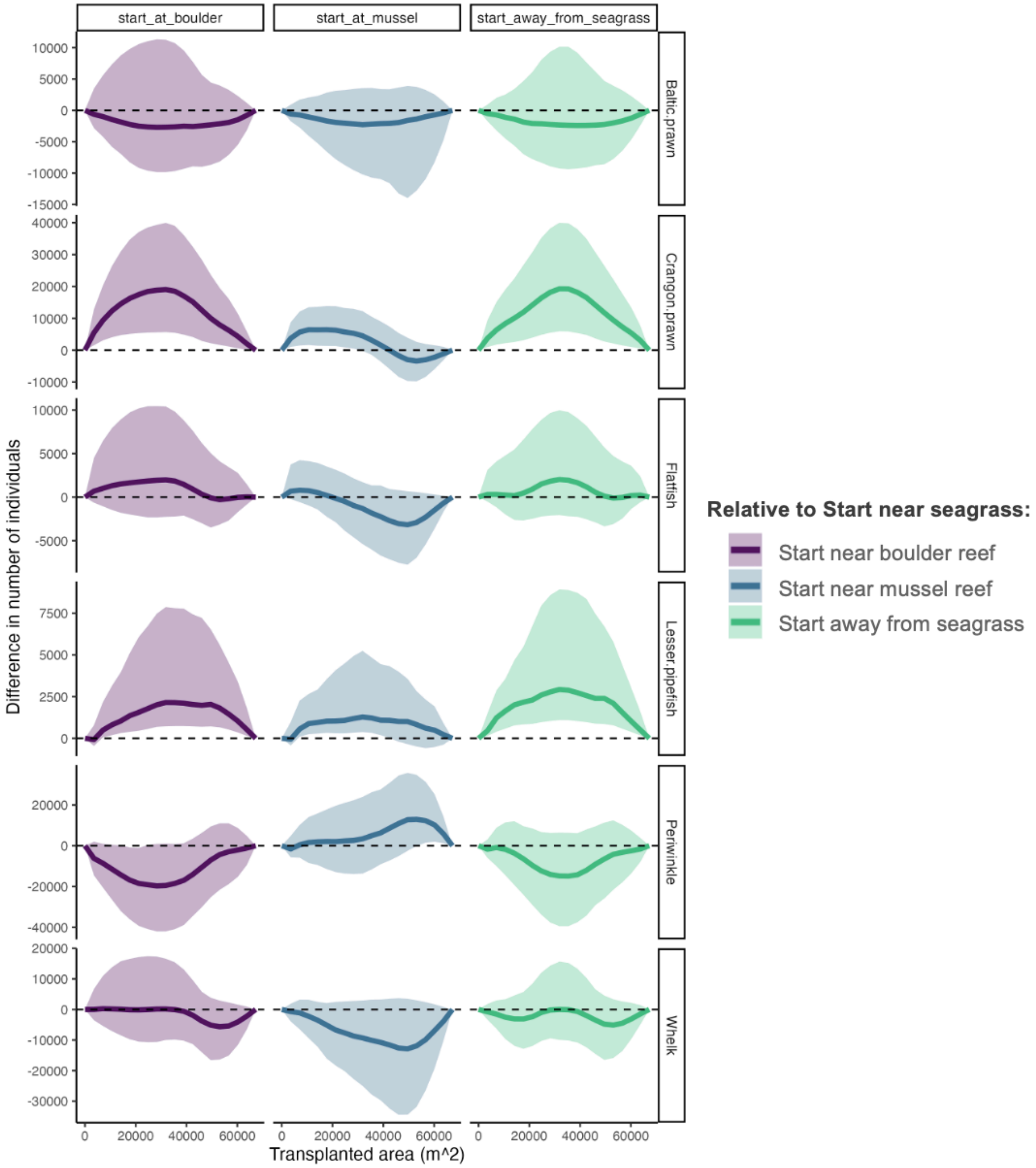


**Fig. S17. Model predictions of differences in the species abundance** relative to the 'start near seagrass' scenario with conditional 95% confidence intervals.

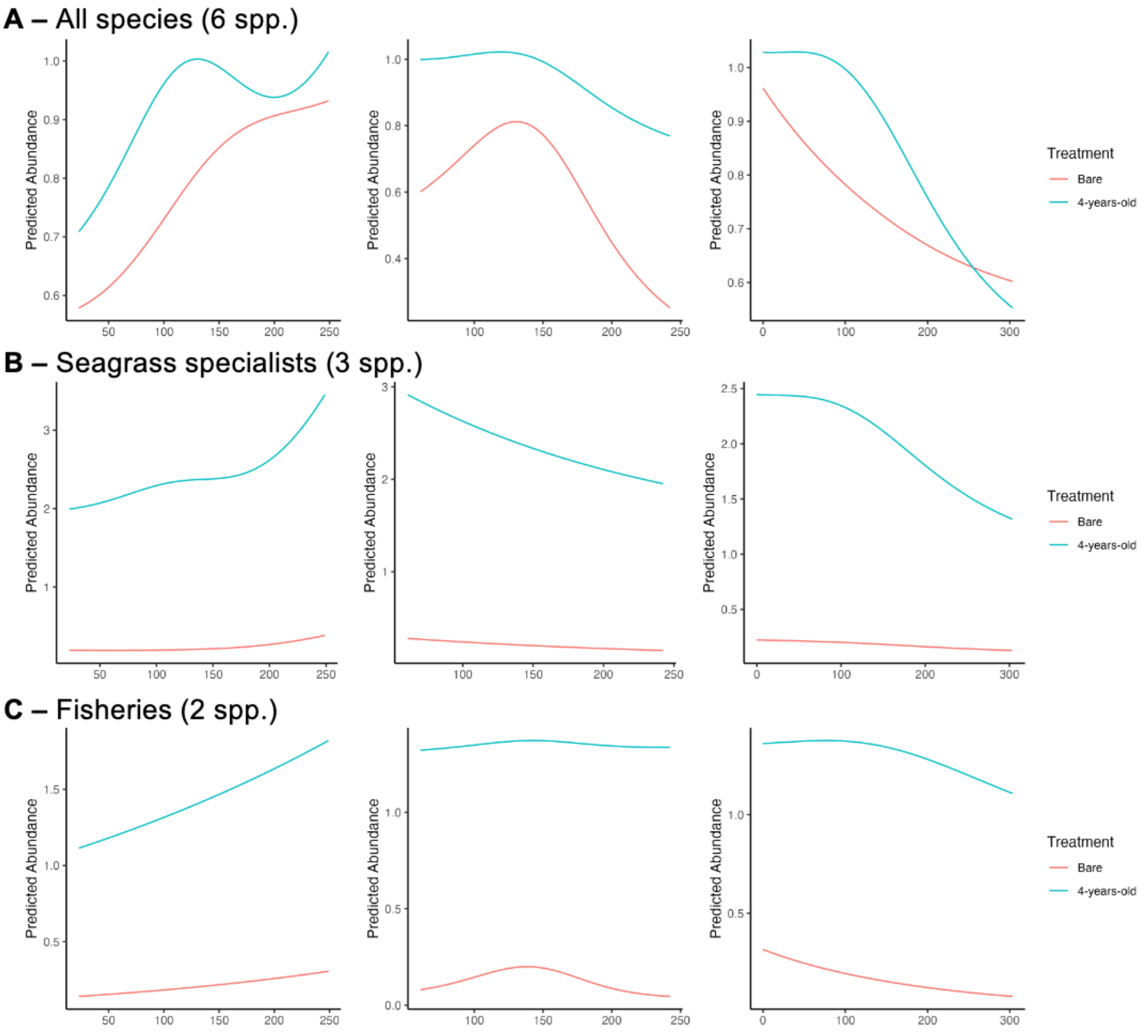


**Fig. S18. Marginal model predictions** (generalised additive models) for the factors: distance to the edge of the boulder reef (DistBouldEdge), distance to the edge of the mussel reef (DistMussEdge), and the area of seagrass in a 10m radius (Area10m) for bare sites (Bare) and restored (4-years-old) sites. Species abundances are scaled by their standard deviations. Abundances are shown for the three mutli-species objective: All species (the six most common species: lesser pipefish (*Sygnathus rostellatus*), common periwinkle (*Littorina littorea*), flatfish (*Pleuronectes platessa* and *Platichthys flesus*), whelks (*Tritia reticulata*), Baltic prawns (*Palaemon adspersus*), and brown shrimp (*Crangon crangon*)); Seagrass specialists (lesser pipefish, common periwinkle, Baltic prawns), and; Fisheries (flatfish, Baltic prawns).

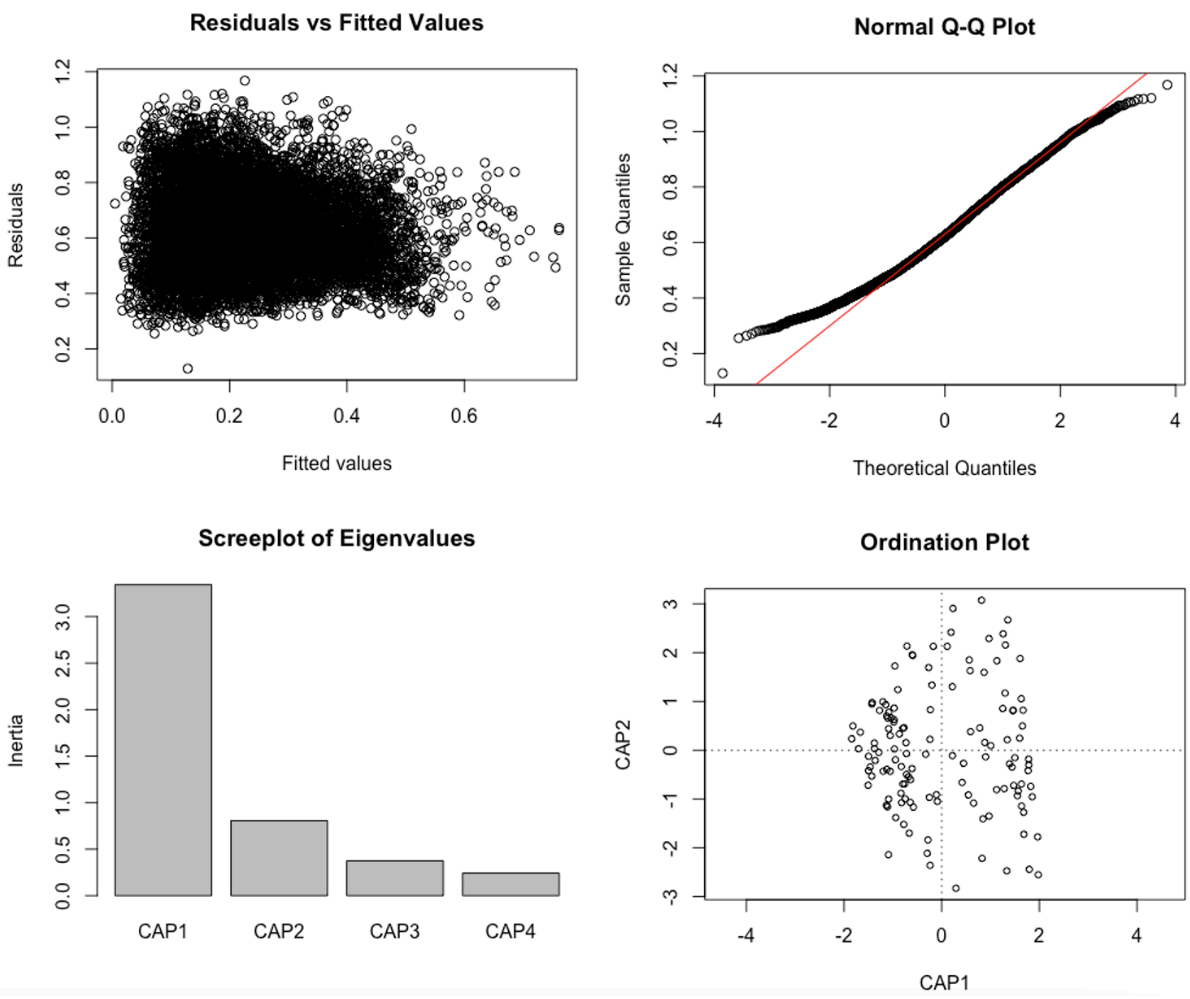


**Fig. S19. Distance-based redundancy analysis (dbRDA) diagnostic plots for beta-diversity**, based on the top 10 most abundant species, a Bray-Curtis dissimilarity matrix, and a permutation test with 999 iterations.

**Table S1. Densities of species (individuals/m$^2$) and their relative frequency (i.e., percentage of sites at which they occurred).**

| Taxon | Common name | Species name | Average density | Relative frequency (%) |
|---|---|---|---|---|
| Crustacean | | | | |
| | Baltic Isopod | *Ideotea balthica* | 0.08 | 15.2 |
| | Baltic prawn | *Palaemon adspersus* | 0.59 | 56.1 |
| | Benthic mysid shrimp | *Praunus flexuosus* | 0.09 | 18.9 |
| | Brown shrimp | *Crangon crangon* | 1.21 | 79.5 |
| | Crab sp. (shore crab) | *Hemigrapsus* sp. | 0.03 | 6.8 |
| | European green crab | *Carcinus maenas* | 0.27 | 49.2 |
| | Hermit crab | *Pagarus bernhardus* | 0.01 | 3.0 |
| | Long legged spider crab | *Macropodia rostrata* | 0.01 | 2.3 |
| | Rockpool shrimp | *Palaemon elegans* | 0.01 | 1.5 |
| Echinoderm | | | | |
| | Brittlestar | *Ophiura albida* | 0.02 | 2.3 |
| | Common starfish | *Asterias rubens* | 0.09 | 20.4 |
| | Green sea urchin | *Psammechinus miliaris* | 0.00 | 0.8 |
| Fish | | | | |
| | Black goby | *Gobius niger* | 0.02 | 6.8 |
| | Lesser pipefish | *Sygnathus rostellatus* | 0.28 | 50.0 |
| | Rock gunnel | *Pholis gunnellus* | 0.00 | 1.5 |
| | Sand goby | *Pomatoschistus minutus* | 0.02 | 6.1 |
| | Sea stickleback | *Spinachia spinachia* | 0.00 | 1.5 |
| | Shorthorn sculpin | *Myoxocephalus scorpius* | 0.01 | 3.3 |
| | Flatfish | *Pleuronectes platessa* and *Platichthys flesus* | 0.09 | 21.2 |
| | Straightnosed pipefish | *Nerophis ophidion* | 0.07 | 18.2 |
| | Three-spined stickleback | *Gasterosteus aculeatus* | 0.00 | 0.8 |
| Gastropod | | | | |
| | Common Periwinkle | *Littorina littorea* | 4.16 | 90.9 |
| | Whelk | *Tritia reticulata* | 1.65 | 84.8 |

**Table S2. Beta diversity of animal communities surveyed within natural seagrass, restored seagrass, and bare sand.** We examined the influence of factors on species composition using distance-based redundancy analysis (dbRDA). Species abundance data for the top 10 most abundant species were used to calculate a Bray-Curtis dissimilarity matrix. We then performed a permutation test with 999 iterations to assess the significance of the predictor variables in explaining variation in species composition.

| Factor | df | SS | F | p |
|---|---|---|---|---|
| **Treatment** | **5** | **8.51** | **9.02** | **0.001** |
| Area of seagrass (10m radius) | 1 | 0.14 | 0.74 | 0.628 |
| Perimeter to area ratio (seagrass within 10m radius) | 1 | 0.19 | 0.99 | 0.381 |
| **Distance to boulder reef** | **1** | **0.48** | **2.53** | **0.017** |
| Distance to mussel reef | 1 | 0.35 | 1.86 | 0.069 |
| Residual | 122 | 23.01 | | |

**Table S3. Spatial independence among community and individual abundance metrics, tested using Moran's I statistic.**

| Response | Observed | p |
|---|---|---|
| Richness | -0.0041 | 0.49 |
| Shannon | -0.0014 | 0.22 |
| Evenness | -0.004 | 0.46 |
| Lesser pipefish | -0.0027 | 0.32 |
| Flatfish | -0.0027 | 0.33 |
| Baltic prawn | -0.0045 | 0.38 |
| Periwinkle | -0.0038 | 0.45 |
| Whelk | -0.0026 | 0.32 |
| Brown shrimp | -0.0028 | 0.34 |

**Table S4. Concurvity between model smoother terms.** Area10m – area of seagrass within a 10m radius; DistBouldEdge – distance to the boulder reef; DistMussEdge - distance to the mussel reef; RatioLog – log-transformed perimeter to area ratio; UTM_E, UTM_N – spatial random effect.

| | s(Area10m) | s(RatioLog) | s(DistBouldEdge) | s(DistMussEdge) | s(UTM_E, UTM_N) |
|---|---|---|---|---|---|
| s(Area10m) | 1 | | | | |
| s(RatioLog) | 0.345 | 1 | | | |
| s(DistBouldEdge) | 0.1729 | 0.0389 | 1 | | |
| s(DistMussEdge) | 0.0959 | 0.0945 | 0.0927 | 1 | |
| s(UTM_E,UTM_N) | 0.5183 | 0.2589 | 0.9219 | 0.6715 | 1 |

**Table S5. ANOVA for reduced 'best' model (based on AIC) for each response variable.** The third column is df for Treatment, and edf for smoother terms. Area10m – area of seagrass within a 10m radius; DistBouldEdge – distance to the boulder reef; DistMussEdge - distance to the mussel reef; RatioLog – log-transformed perimeter to area ratio; UTM_E, UTM_N – spatial random effect.

| Response | Factor | edf/df | Ref.df | Chi.sq | p |
|---|---|---|---|---|---|
| Richness | **Treatment** | **5** | | **7.22** | **<0.001** |
| | **s(DistMussEdge)** | **1** | **1** | **12.25** | **0.001** |
| Shannon | s(Area10m) | 1.743 | 1.933 | 3.77 | 0.051 |
| | s(DistBouldEdge) | 1 | 1 | 2.84 | 0.094 |
| | **s(DistMussEdge)** | **1** | **1** | **9.68** | **0.002** |
| | s(UTM_E,UTM_N) | 2.876 | 2.984 | 2.59 | 0.064 |
| Evenness | **Treatment** | **5** | | **4.18** | **0.002** |
| | s(DistMussEdge) | 1 | 1 | 3.10 | 0.081 |
| Lesser | **Treatment** | **5** | | **22.11** | **<0.001** |
| | **s(Area10m)** | **1.822** | **1.968** | **6.83** | **0.022** |
| | **s(DistBouldEdge)** | **1.917** | **1.993** | **9.96** | **0.006** |
| Flatfish | **Treatment** | **5** | | **18.32** | **0.003** |
| | **s(DistMussEdge)** | **1.884** | **1.986** | **6.67** | **0.030** |
| | s(RatioLog) | 1 | 1 | 3.58 | 0.058 |
| Baltic prawn | s(DistBouldEdge) | 1 | 1 | 2.89 | 0.089 |
| | **s(RatioLog)** | **1.896** | **1.989** | **8.32** | **0.015** |
| Periwinkle | **Treatment** | **5** | | **132.30** | **<0.001** |
| | s(DistBouldEdge) | 1.761 | 1.943 | 4.45 | 0.072 |
| | **s(DistMussEdge)** | **1** | **1.001** | **4.33** | **0.038** |
| Whelk | s(Area10m) | 1 | 1 | 3.84 | 0.050 |
| | **s(UTM_E,UTM_N)** | **2** | **2** | **12.39** | **0.002** |
| Brown shrimp | **Treatment** | **5** | | **24.64** | **<0.001** |
| | s(DistBouldEdge) | 1 | 1 | 3.38 | 0.066 |
| | **s(DistMussEdge)** | **1.691** | **1.905** | **12.14** | **0.001** |
| | **s(UTM_E,UTM_N)** | **2.881** | **2.986** | **16.71** | **0.002** |
